# The Cul5 E3 Ligase Complex Is a Key Negative Feedback Regulator of TCR/IL2 Signaling and Anti-Tumor Activity in CD8^+^ T Cells

**DOI:** 10.1101/2022.11.16.516824

**Authors:** Xiaofeng Liao, Wenxue Li, Ao Li, Barani Kumar Rajendran, Jingjing Ren, Hongyue Zhou, David Calderwood, Benjamin Turk, Wenwen Tang, Yansheng Liu, Dianqing Wu

## Abstract

CD8^+^ T cells play an important role in tumor immune surveillance and control. Better understanding of the regulation of their anti-tumor actions and improving their cytotoxic function and persistence will help advancing cancer immunotherapies. Here, we report the development of a step-wise CRISPR knockout (KO) screening strategy under the selection of TGF-β, a clinically relevant immunosuppressive pressure. The screen identifies Cul5 as a negative-feedback regulator of the core signaling pathways, differentiation, and persistence of CD8^+^ T cell. Cul5 KO in mouse CD8^+^ T cells significantly improves their tumor control ability *in vitro* and *in vivo* with significant proteomic alterations that generally enhance TCR and cytokine signaling, effector function, stemness, and survival of CD8^+^ T cell. Mechanistically, Cul5, whose protein content and active, neddylated form increase upon TCR-stimulation, interacts with SOCS-box-containing Pcmtd2 and negatively regulates TCR and IL2/STAT5 signaling by decreasing TCR and IL2 signaling molecules. Moreover, Cul5 KO in human CD8^+^ T cells phenocopies that in mouse CD8^+^ T cells. Furthermore, KO of CTLA4 that is markedly upregulated by Cul5 KO in mouse and human CD8^+^ cells further enhances anti-tumor effect of Cul5 KO, and a neddylation inhibitor enhances CD8 effector activities largely dependently of Cul5. These results together not only reveal a previously unknown negative-feedback regulatory mechanism for CD8^+^ T cells, but also have strong translational implications in cancer immunotherapy.

## Main

CD8^+^ T cells play a central role in cancer immunotherapy including immune checkpoint inhibition (ICI) and adaptive cell transfer (ACT). Current approved ICI in clinic relies on the re-activation of anti-tumor CD8^+^ T cells by neutralizing co-inhibitory molecules CTLA4 and/or PD1/PD-L1^1–4^. Antibodies targeting other co-inhibitory and co-stimulatory molecules such as Tim3, Lag3 and 4-1BB are being actively developed and investigated in clinical trials to further release the power of anti-tumor CD8^+^ T cells^5, 6^. ACT including tumor infiltrating lymphocytes (TILs)^7–10^, chimeric antigen receptor engineered T (CAR-T) cells^11–14^ and T cell receptor engineered T (TCR-T) cells^15–17^ also has shown promising clinical efficacy in a subset of cancer patients with malignancies otherwise refractory to other treatments. However, many hurdles still exist to prevent the successful application of ICI and ACT to the remaining majority of patients^18, 19^. The immunosuppressive tumor microenvironment (TME) is one of the critical hurdles as it diminishes the effector functions and persistence of pre-existing endogenous anti-tumor CD8^+^ T cells as well as adoptively transferred T cells^20^. Genome-wide screens in T cells have been performed to identify genes that either positively or negatively regulate anti-tumor functions of cytotoxic T cells^21–27^, providing new opportunities to overcome the immunosuppressive hurdles in tumors. However, as enrichment or depletion-based screening systems are highly context-dependent, the selection pressures and criteria applied in different studies will result in the identification of differential targets, suggesting that proper selection of a clinically relevant screening pressure should result in better translation into clinical application. Among various immunosuppressive factors in TME, Transforming Growth Factor (TGF)-β appears to be a common one that directly suppresses T cell proliferation and effector functions, while promoting the exhaustion of cytotoxic T cells^28–30^. Although directly targeting TGF-β proximal signaling by overexpression of dominant negative TGFbRII on CAR-T cells revealed a promising efficacy and safety in a recent reported phase I clinical trial, TGF-β as a pleiotropic cytokine also maintains an immune homeostasis to prevent autoimmunity^31^, facilitates the formation of resident memory^32–34^ and memory precursor T cells^30^, and prevents the malignant transformation of pre-malignant cells^35^. Therefore, it is clinically important to identify other proteins either in the distal TGF-β signaling pathway or in parallel pathways that can counteract TGF-β immunosuppressive effects without systemically severe inflammatory side effects, compromising the persistence, or possibly increasing malignant transformation of adoptively transferred cytotoxic T cells.

E3 ubiquitin ligases have been shown to regulate T cell responses via the ubiquitination and subsequent degradation of TCR signaling molecules^36^. Cul5 is a key scaffold molecule in the cullin-ring E3 ligase (CRL) complex that consists of Elongin C, Elongin B, Rbx2, and one of the SOCS-box-containing proteins. The SOCS-box-containing proteins act as the receptors for a specific set of substrate proteins and mediate their ubiquitination and degradation^37^. The CIS/SOCS family SOCS-box-containing proteins bind to Cul5 with variable affinities through the SOCS-box domain^38^, and some of these proteins including Cish, SOCS1 and SOCS3 have been shown to play regulatory roles in T cell activation by targeting cytokine and TCR signaling^39, 40^. However, it is not known whether the Cul5 E3 ligase complexes regulate CD8^+^ T cell activation or function through any of these CIS/SOCS family proteins in CD8^+^ T cells despite Cul5 regulates pJak1 degradation by interacting with Cish in CD4 cells^41^. Neddylation is a critical modification of CRLs to induce their conformational changes and subsequent activation^42^. As overactivated neddylation of CRLs is associated to disease progression and poor survival of multiple human cancers^43–45^, neddylation inhibitors blocking E1 NEDD8-activating enzyme such as MLN4924 and TAS4664 have been developed and investigated in over 40 clinical trials (https://www.clinicaltrials.gov/) to evaluate their safety and anti-tumor efficacy. Besides its effects on tumor cells, neddylation has been shown to regulate the functions of various immune cells in TME including CD4^+^ T cells^46, 47^. However, little is known about the effect of neddylation on CD8^+^ T cells.

In this study, we developed a step-wise CRISPR screening approach by performing whole-genome *in vitro* screens in the presence of TGF-β and targeted *in vivo* bulk screens, followed by *in vivo* single-cell screens. This approach allowed us to robustly identify gene targets, including Cul5, in CD8^+^ T cells that improved anti-tumor efficacy under TGF-β-induced immunosuppressive pressure. Proteomic analyses further revealed an understudied SOCS-box-containing protein, Pcmtd2, as the dominant substrate receptor of the Cul5 E3 ligase complex, which negatively regulates TCR and IL2/STAT5 signaling by targeting TCR and IL2 signaling molecules including the TCR/CD3 complex and IL2 receptor β subunits (Il2rb).

### Step-wise bulk CRISPR KO screens enrich potential ACT boosters

To identify TGF-β resistant targets, we developed a step-wise CRISPR screening approach combining an initial T cell line-based *in vitro* genome-wide screening with primary CD8^+^ T cell *in vivo* screens at the bulk and then single cell levels (Fig. 1a). This design allowed us to circumvent the cost and labor ineffectiveness of performing a genome-wide *in vivo* CRISPR/Cas9 screen, which, due to library size and the very limited number of tumor-infiltrating T cells^21, 24^, would require a large number of mice. IL2 is important for T cell proliferation and function *in vivo* ^48^ and for generation of adoptive transferred T cell products *in vitro*^49^, while TGF-β is a common and dominant immunosuppressive factor in TME^50^. Therefore, for the initial genome-wide CRISPR screen, we used an immortal mouse T cell line, HT2, whose proliferation is IL2 dependent^51^ and suppressed by TGF-β^52^. The HT2 T cells were transduced with the Briea genome-wide mouse CRISPR KO library in a lentiviral vector co-expressing spCas9 and sgRNA ^53^. As shown in Extended Data Fig. 1b, compared to IL2 culture alone, the addition of TGF-β1 significantly but not completely suppressed the proliferation of CRISPR library transduced HT2 cells. We used the following three criteria to select putative candidate hit sgRNAs: 1) they should not interrupt IL2-dependent cell survival and proliferation, which means sgRNAs depleted in Day-21 IL2 culture alone compared to pre-culture input should be excluded from consideration; 2) They should resist TGF-β1-dependent suppression of proliferation, which means sgRNAs enriched in Day-21 IL2+TGF-β1 culture compared to Day-21 IL2 culture alone should be included in consideration; 3) the candidate genes should have detectable transcription levels in primary CD8^+^ T cells either at resting state or after activation. With these selection criteria, we set a sgRNA enrichment (Log_2_ FC of IL2+TGF/IL2) cutoff > 1.585 for candidate inclusion while a sgRNA depletion (Log_2_FC of IL2/input) cutoff < -2 for exclusion (Fig. 1b). We found that sgRNA hits for several known TGF-β signaling molecules or inducible genes were enriched, including Tgfbr1, Tgfbr2, Smad2, Smad4, Cxxc5, Cdkn2a, and Ahnak^50, 54–56^, while sgRNAs targeting known essential genes for T cell survival and IL2-dependent activation were significantly depleted, including Il2ra, Stat5a, Akt1, Mapk1, and Jak3^48^. These results provide a validation for this *in vitro* genome-wide screening. After further excluding the genes already known related to TGF-β or not detectable in primary mouse CD8^+^ T cells by Immgen dataset ^57^, 270 candidate genes were selected for further *in vivo* screens in primary CD8^+^ T cells (Supplementary Table 1).

**Fig. 1:**
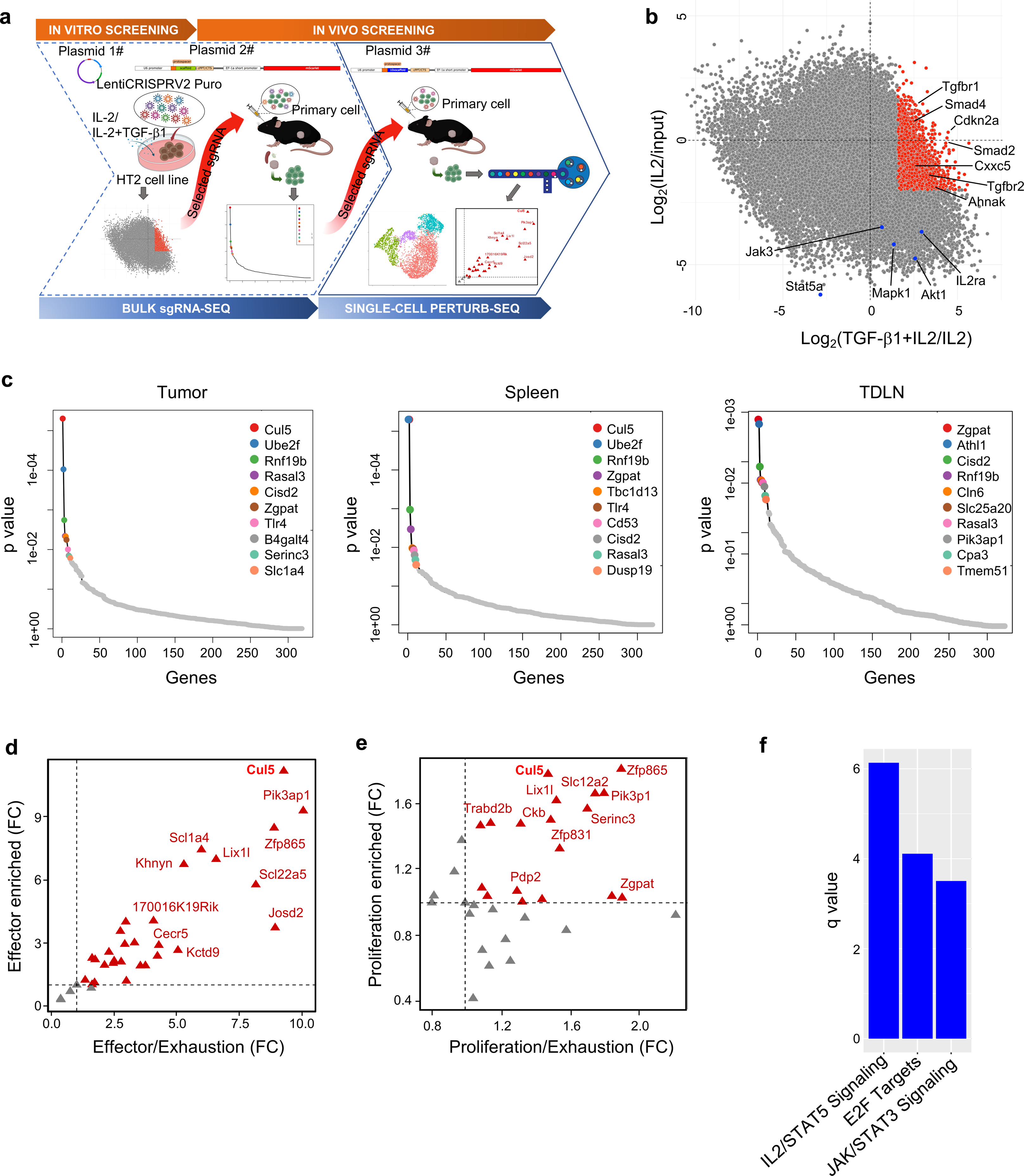
Step-wise CRISPR KO screens identify genes enhancing anti-tumor activity of CD8^+^ T cells. **a,** Schematic representation of the experimental design for step-wise CRISPR KO screens. **b,** Scatter plots of sgRNA log_2_-fold change (x-axis, TGF-β1+IL2/IL2; y-axis, IL2/input) in HT2 cells cultured *in vitro* for 21 days. **c,** Gene enrichment analysis of the bulk *in vivo* screen in tumors (Left), spleens (Middle), and TDLNs (Right) using the MAGeCK analysis. **d,e,** Genes enriched in the **d,** effector cluster and **e,** proliferating cluster in the single cell CRISPR KO screening are presented as the target gene enrichment over the non-targeting control being plotted against the ratio of the enrichment in **d,** effector cluster or **e,** proliferating cluster to that in the exhausted cluster. **f,** Signaling pathways enriched in transferred tumor-infiltrating Cas9/OT-I cells expressing Cul5 sgRNAs compared to those expressing non-targeting sgRNAs.

We used OT-I T cells against ovalbumin (OVA)-expressing tumor as an *in vivo* ACT model to perform the secondary CRISPR screening. We compared TGF-β expression levels among four commonly used C57Bl mouse syngeneic cancer cell lines and found that EL4 expressed the highest level of TGF-β (Extended Data Fig. 1c). Thus, we chose the corresponding OVA-expressing EL4 cell line E.G7-OVA for further screens to ensure the presence of enhanced TGF-β immunosuppressive microenvironment. We generated Cas9-expressing OT-I (Cas9/OT-I) T cells by crossing OT-I and Cas9 mice. Cas9 and OT-I TCR expression in CD8^+^ T cells were confirmed by flow cytometry (Extended Data Fig. 1d), and OVA antigen specific T cell activation was validated by degranulation and cytokine production after co-culture with E.G7-OVA cells (Extended Data Fig. 1e). Gene knockout efficiency over 70% of Cas9/OT-I T cells was confirmed with CD8a KO by retroviral transduction of an sgRNA targeting CD8a (Extended Data Fig. 1f). A customized sgRNA library containing sgRNAs (six for each target gene) targeting the 270 selected genes was generated with the coverage and normal distribution of sgRNAs confirmed by next generation sequencing (Extended Data Fig. 1g). Library-transduced Cas9/OT-I T cells were sorted based on the mScarlet reporter expression and then transferred into E.G7-OVA tumor-bearing mice. Upon T cell transfer, tumors shrunk rapidly in the first 7 days and then became relatively stable for the next 5 days (Extended Data Fig. 1h), suggesting a switch of T cell anti-tumor immune response from an active effector stage to an immune suppression/exhaustion stage as previously reported ^58^. Because tumors can also suppress immune responses systemically^59, 60^, library-transduced T cells were collected from tumors, tumor-draining lymph nodes (TDLNs), and spleens by sorting CD8^+^GFP^+^ cells at day 12 post T cell transfer. We reasoned that the proliferation and persistence of immune suppression-resistant T cells should outcompete others, which can be reflected by the enrichment of sgRNAs in the library. Therefore, sgRNA enrichment analysis was carried out by comparing each sample with the pre-transfer input. As the step-wise screening significantly limited *in vivo* sgRNA pool, an average of 150x coverage per sgRNA per tissue was achieved from recovered T cells of one recipient. A total of 4 replicates for each tissue were then used for robust sgRNA enrichment analysis. Fig. 1c showed top enriched gene hits from the tumor, spleen and TDLN, respectively. Cul5, Ube2f and Rnf19b were top three hits in both tumor and spleen, while Zgpat, Athl1 and Cisd2 were top three hits in TDLN. Interestingly, Cul5 and Ube2f are the components of the Cul5-E3 ligase complex, suggesting a possible important role of the E3 ligase complex in the regulation of CD8^+^ T cell anti-tumor responses.

### Single cell CRISPR screening identifies Cul5 as a CD8 cell regulator

The above two CRISPR screens are based on an overall T cell number enrichment, which reflects the proliferation and persistence of T cells as an entire population. However, tumor-infiltrating T cells are known to have a variety of differentiation status and subpopulations^61, 62^, complicating the interpretation of the bulk CRISPR screen results. Moreover, a superior anti-tumor cytotoxic T cell response also requires enhanced cytotoxic effector functions besides the increase of total cell number^21, 25^. Previous effector based CRISPR screens are biased to a single phenotype such as degranulation (surface CD107a^+^)^21^ or cytokine production (IL2 or IFNg intracellular staining)^63^ which may not be able to completely reflect functional T cell subsets. Therefore, to comprehensively identify gene candidates regulating functional CD8^+^ T cell subgroups, we decided to perform single-cell CRISPR KO screening with tumor-infiltrating CD8^+^ T cells by single-cell RNA sequencing in parallel with single-cell sgRNA sequencing. To make our CRISPR system compatible with 10x genomics kit, the original sgRNA scaffold sequence was replaced by the one containing 10x genomics capture sequence in the stem loop region. Gene editing efficiency as high as over 80% with the new scaffold replacement was confirmed by CD8a KO in primary Cas9/OT-I T cells (Extended Data Fig. 2a). Thirty-three candidate genes with 3 top efficient sgRNAs per gene plus 4 non-targeting sgRNAs were selected from the *in vivo* bulk screening with a selection criteria of more than one sgRNA enrichment over 1.4 fold per gene in each sample (Supplementary Table 2). Seven days post ACT in the E.G7-OVA model, over 10,000 sorted tumor-infiltrating Cas9/OT-I T cells were subjected to 10x single-cell RNA sequencing to obtain approximate 300x coverage per target gene. Unsupervised clustering divided the cells into four clusters that were visualized in 2D Uniform Manifold Approximation and Projection (UMAP) (Extended Data Fig. 2d). Among differential expressed gene (DEG) set for each cluster (Extended Data Fig. 2b), T cell phenotype markers were used to annotate the four clusters into exhausted-like cells (Ccl3, Ccl4, Tnfrsf9, Ifng, Lag3 and Harvcr2) as Cluster 0; proliferating cells (Mik67 and Cdk1) as Cluster 1; resting precursor-like cells (Il7r, S1pr1, Tcf7 and Slamf6) as Cluster 2; effector cells (Gzmc, Gzmd, Gzme, Gzmf, Prf1, Spp1, Klre1 and Klrd1) as Cluster 3 (Extended Data Fig. 2c)^62^. UMAP cell cycle phase analysis also showed that the majority of the cells in the G2M and S stages were in cluster 1, whereas the other three clusters were mainly in the G1 stage (Extended Data Fig. 2e), consistent with the proliferating cell identity for Cluster 1. As the exhaustion-like cluster is the unfavorable group in anti-tumor responses of cytotoxic T cells, we evaluated the enrichment of individual gene KO T cells in the other three functional clusters based on both the absolute cell counts and respective ratio to the exhaustion cluster, with the pre-transfer input and non-targeting sgRNA transduced T cells as the normalization control (Fig. 1d,e and Extended Data Fig. 2f). This systemic and unbiased functional analysis of the screen revealed that Cul5 KO cells were highly enriched in both effector and proliferating clusters. Several other hits, such as Pik3ap1, Zfp865 and Lix1l, were also enriched in two or three of these functional clusters.

### Cul5 KO enhances anti-tumor effects of primary CD8^+^ T cells in vivo

In the above single cell analysis, the effector cluster showed much higher enrichment magnitude compared to the proliferating and precursor-like clusters (Fig. 1d,e and Extended Data Fig. 2f), suggesting the stronger impact of gene perturbation on the effector population. Among top enriched gene (Pik3ap1, Cul5 and Zf865) KO in the effector cluster, the role of Pik3ap1 in CD8^+^ T cell activation through PI3K signaling has been reported^64^, while Zf865 is a mouse specific gene. Cul5 as a core element of E3 ligase complexes, on the other hand, has not been investigated in CD8^+^ T cells, despite a recent study reporting its regulation of CD4^+^ T helper cell differentiation^41^. Therefore, we decided to investigate Cul5 further. The fact that the Cul5 gene expression level was much lower in the Cul5 KO cells than non-targeting KO cells or other gene KO cells (Extended Data Fig. 2g) confirms that the Cul5 sgRNAs yielded a high KO efficiency. A further gene enrichment and signaling pathway analysis of DEG between the Cul5 KO and non-targeting KO cells indicated that the IL2/STAT5, E2F targets and JAK/STAT3 signaling pathways were significantly altered (Fig. 1f). Taken together, our step-wise CRISPR KO screens revealed Cul5 may play an important role in regulating the persistence and effector functions of tumor-reactive CD8^+^ T cells.

To validate the enhanced anti-tumor responses of Cul5 KO CD8^+^ T cells, Cas9/OT-I T cells transduced with the Cul5 sgRNAs or a non-targeting control sgRNA were adoptively transferred into E.G7-OVA tumor-bearing mice subjected to prior sub-lethal irradiation for lymphodepletion^65^. The Cul5 KO T cells showed significantly stronger tumor control ability than the NC T cells (Fig. 2a), resulting in significantly improved survival (Fig. 2b). Further analysis of tumor-infiltrating transferred T cells as well as those in TDLNs at the end point revealed that the Cul5 KO T cells had significantly higher IL7R (Fig. 2c and Extended Data Fig. 3a) and GZMB (Fig. 2d and Extended Data Fig. 3b) expression levels than the NC cells. These results suggest that the Cul5 KO T cells possess a hybrid stemness/effector phenotype, which may explain their improved functional persistence. Concordantly, the Cul5 KO group had a significantly higher number of tumor-infiltrating transferred T cells that were still capable of producing IFNg and/or TNF upon *in vitro* re-stimulation than the NC group (Fig. 2e and Extended Data Fig. 3c). In addition, transfer of the Cul5 KO T cells reduced the incidence of TDLN-metastasis of primary E.G7-OVA tumors from 65% to 20% (Fig. 2f), suggesting that the Cul5 KO T cells may suppress metastasis along with a better control of primary tumors. TDLN has been demonstrated as a reservoir of tumor-specific stem-like CD8^+^ T cells^66^ that preserve the ability of multiple cytokine-producing ability upon re-stimulation^67^ and are important for ICI efficacy and sustained anti-tumor responses^68^. In this study, we observed a negative correlation of the proportion of transferred stem-like CD8^+^ T cells in TDLN capable of producing multiple cytokines with TDLN metastasis (Fig. 2g) regardless of Cul5 KO. This observation suggests that established metastatic tumor cells may induce exhaustion of anti-tumor stem-like CD8^+^ T cells in TDLN, which is consistent with a similar finding in a clinical breast cancer metastasis study^69^. Concordantly to the reduced metastasis incidence, we also observed an increased incidence of TDLN possessing high proportion of stem-like CD8^+^ T cells (multiple cytokine producer upon restimulation, from 40% to 80%) in the Cul5 KO group (Fig. 2h), suggesting a better preservation of functional stem-like CD8^+^ cells with Cul5 KO. These results together indicate that Cul5 KO in T cells have an increased hybrid stemness/effector phenotype that enhances their anti-tumor effects in both primary tumor suppression and metastasis prevention *in vivo*.

**Fig. 2:**
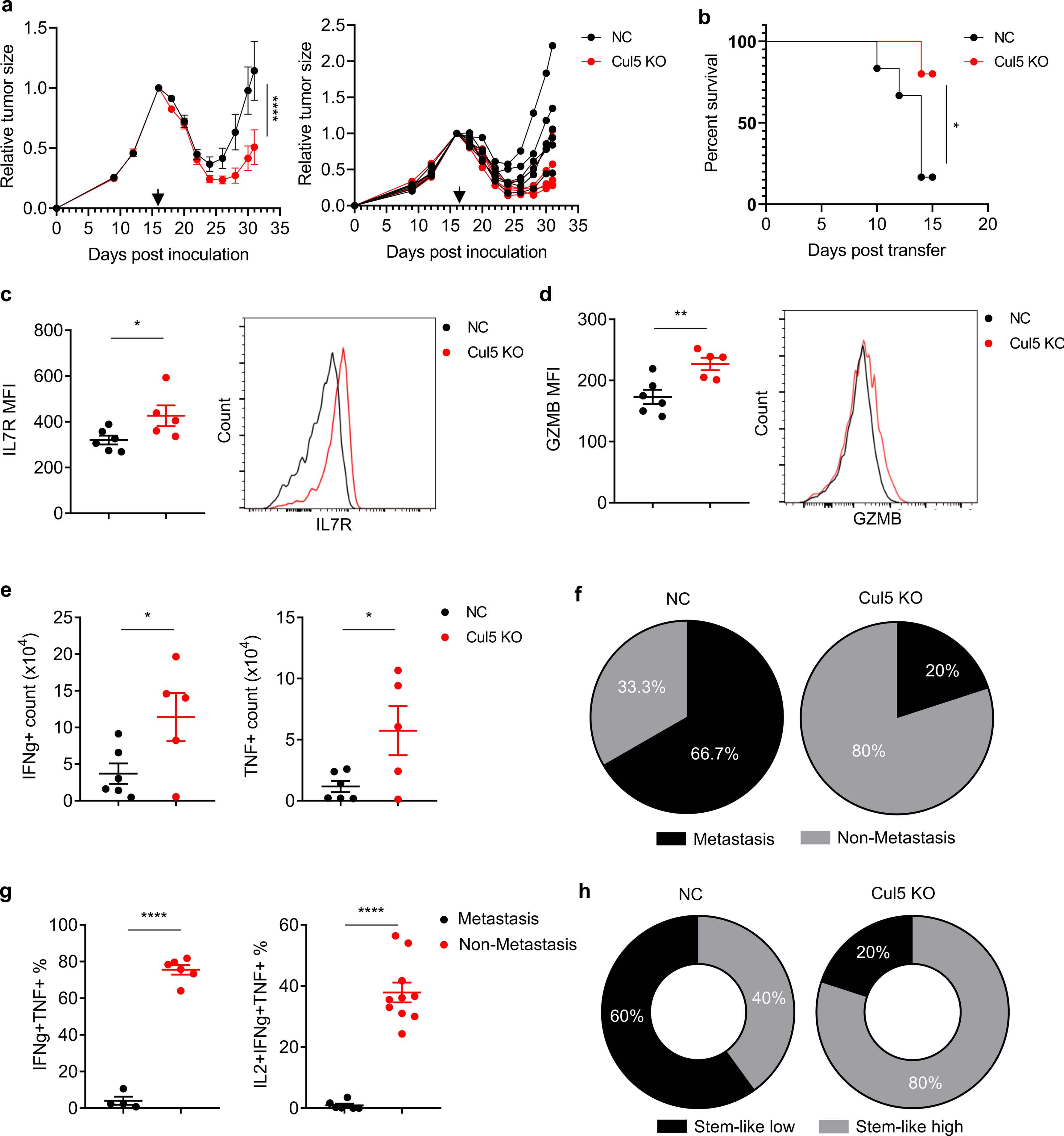
Cul5 KO in primary CD8^+^ T cells enhanced anti-tumor responses. **a,** The effect of the transfer of Cul5 KO or non-targeting (NC) OT-I CD8+ T cell on tumor progression was examined in the C57BL/6N mice inoculated with the E.G7-OVA cells. Data are shown as mean ± SEM on the left and as individual mouse on the right. Black arrow indicates the time of sub-lethal irradiation followed by immediate adoptive transfer of Cul5 (Red) KO or NC (Black) Cas9/OT-I cells. ****p<0.0001 by two-way ANOVA (n=5-6 mice per group). **b,** Survival curve of E.G7-OVA tumor-bearing C57BL/6N mice post adoptive transfer of Cul5 KO (Red) or NC (Black) Cas9/OT-I cells. *p<0.05 by Gehan-Breslow-Wilcoxon test (n=5-6 mice per group). **c,d,** Flow cytometry analysis of **c,** CD127 and **d,** Granzyme B expression of transferred tumor-infiltrating Cul5 KO or NC Cas9/OT-I cells. Representative histograms are shown. Data are shown as mean ± SEM (*p < 0.05 and **p < 0.01, by unpaired t-test). **e,** Flow cytometry analysis of IFNg^+^ (Left) and TNF^+^ (Right) cell counts per tumor of transferred tumor-infiltrating Cas9/OT-I cells post re-stimulation *in vitro*. Data are shown as mean ± SEM (*p < 0.05, by unpaired t-test). **f,** Pie chart showing the percentage of lymph node (LN) metastasis (Black) and non-metastasis (Grey) in each group. **g,** Flow cytometry analysis of IFNg^+^TNF^+^ (Left) or IL2^+^IFNg^+^TNF^+^ (Right) transferred TDLN-infiltrating Cas9/OT-I cells post re-stimulation *in vitro* in LNs with (Black) and without metastasis (Red). Data are shown as mean ± SEM (****p < 0.0001, by unpaired t-test). **h,** Donut chart showing the percentage of exhaustion subtype (Black)-dominant and progenitor subtype (Grey)-dominant transferred TDLN-infiltrating Cas9/OT-I cells in each group.

### Cul5 KO enhances effector activities of primary CD8^+^ T cells *in vitro*

To further investigate the effects of Cul5 KO on primary CD8 T cell responses, we generated Cul5 KO CD8^+^ T cells using cells isolated from the spleens of the Cas9/OT-I mice, and performed cytokine-induced differentiation, TCR-dependent activation, and tumor cell killing assays *in vitro*. Continued differentiation and expansion of Cul5 KO T cells in the IL2, IL7 and IL15 cytokine cocktail prior to TCR-dependent stimulation already showed increased intracellular GZMB and IFNg expression compared to the NC T cells (Fig. 3a), suggesting that Cul5 is actively involved in cytokine-dependent cytotoxic T cell differentiation. We then examined multiple effector markers using flow cytometry 6 and 16 hours post anti-CD3 stimulation in the presence or absence of TGF-β1. GZMB and IFNg were significantly higher at both time points in the Cul5 KO cells than the NC ones (Fig. 3b,c,d,e). In agreement with flow cytometry results, secreted IFNg detected by ELISA was significantly higher in Cul5 KO cells than the NC cells as well (Fig. 3f). GZMB expression at 16 hours and IFNg expression at both 6 hours and 16 hours were significantly suppressed by TGF-β1 to similar degrees in Cul5 KO and NC cells (Fig. 3c,d,e), suggesting that Cul5 may be not directly involved into TGF-β signaling and cannot be completely suppressed by TGF-β. Because Gzmb expression (Fig. 3b,d) and IFNg production (Fig. 3e,f) in the Cul5 KO cells in the presence of TGFb1 were still significantly higher than those in its absence, Cul5 KO would present an apparent TGFb1 resistance. When co-cultured with the E.G7-OVA cells, *in vitro* differentiated OT-I T cells with Cul5 KO showed significantly higher tumor cell killing ability at several non-saturated effector to target ratios than the NC cells (Fig. 3g), accompanied with significantly increased GZMB and IFNg expression levels upon E.G7-OVA cell stimulation over those of the NC cells (Fig. 3h). These data together connect Cul5 to the regulation of cytotoxic T cell differentiation, activation, and effector functions.

**Fig. 3:**
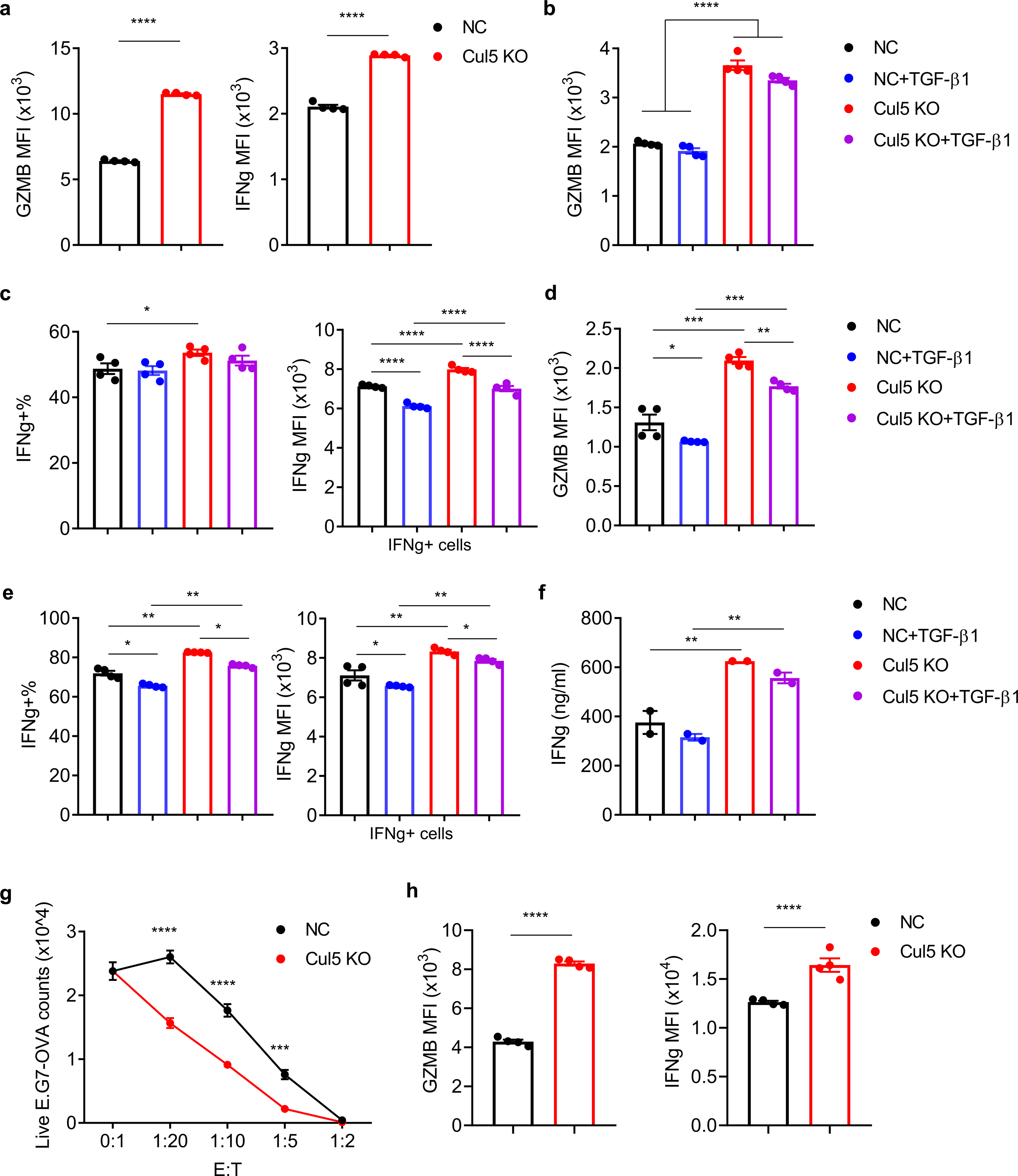
Immunological characterization of Cul5 KO primary CD8^+^ T cells *in vitro*. **a-e,** Flow cytometry analysis of effector molecules (GZMB and IFNg) in NC (Black and Blue) or Cul5 KO (Red and Purple) primary CD8^+^ T cells **a,** prior to or post anti-CD3+anti-CD28 stimulation for **b,c,** 6 hours or **d,e,** 16 hours in the presence (Blue and Purple) or absence (Black and Red) TGF-β1. Data are shown as mean ± SEM (*p < 0.05, **p < 0.01, ***p < 0.001, and ****p < 0.0001 by unpaired t-test, n=4). **f,** ELISA analysis of secreted IFNg from NC (Black and Blue) or Cul5 KO (Red and Purple) primary CD8^+^ T cells 16 hours post anti-CD3+anti-CD28 stimulation in the presence (Blue and Purple) or absence (Black and Red) TGF-β1. Data are shown as mean ± SEM (**p < 0.01 by unpaired t-test, n=2). **g,** *In vitro* cytotoxicity assay of NC (Black) or Cul5 (Red) KO OT-I cells co-cultured with E.G7-OVA cells overnight. E:T, effector (T cell) to target (tumor cell) ratios. Data are shown as mean ± SEM (***p < 0.001 and ****p < 0.0001 by multiple t-test, n=4). **h,** Flow cytometry analysis of GZMB and IFNg expression of NC (Black) or Cul5 KO (Red) OT-I cells co-cultured with E.G7-OVA cells at a E:T ratio of 1:20 for 6 hours. Data are shown as mean ± SEM (****p < 0.0001 by unpaired t-test, n=4).

### Cul5 KO alters CD8^+^ T cell proteome

To investigate the mechanism by which Cul5 regulates CD8^+^ T cell cytokine-dependent differentiation and TCR-dependent activation, we performed quantitative mass spectrometry (MS) by data-independent acquisition (DIA)-MS ^70^ of total proteins in the Cul5 KO and NC primary CD8^+^ T cells in the following conditions: 1) Cytokine dependent expansion and differentiation (T0); 2) 8 hours cytokine withdraw prior to TCR stimulation (T8); and 3) 16 hours TCR stimulation post 8-hour cytokine withdraw (T16). Principal component analysis (PCA) and correlation analysis revealed that replicates in each condition clustered together while different conditions separated from each other (Extended Data Fig. 4a,b), suggesting significant proteomic changes among different conditions of the same T cells as well as between Cul5 KO and NC T cells in each condition. Together with similar total protein quantities among all detected samples (Extended Data Fig. 4c), DIA-MS analyses were of high quality. Additionally, the strong reductions of Cul5 abundances in the Cul5 KO cells from all three conditions compared to the NC cells (Fig. 4a) confirms high KO efficiency. Of note, the proteomic analysis showed that the Cul5 protein level was significantly increased upon TCR stimulation post cytokine starvation in the NC cells, suggesting a potential negative feedback regulatory role of Cul5 in CD8^+^ T cell activation (Fig. 4a). Consistent with this idea, we observed more markedly upregulation of Cul5 expression upon of TCR stimulation of naïve primary CD8^+^ cells than TCR restimulation (Fig. 4b). More importantly, TCR stimulation also increased neddylation^42, 71^ of Cul5 in these primary CD8^+^ T cells (Fig. 4c). Upon differential expression analysis of the proteomic data with stringent cutoffs (p value<0.01 and FC >1.8 or <0.55), we found 184 and 69 in T0; 177 and 65 in T8; 234 and 79 in T16 up- and down-regulated proteins, respectively, in the Cul5 KO T cells compared to NC T cells (Supplementary Table 3 and Fig. 4d). The commonly and uniquely regulated proteins among three conditions are shown in the Venn diagram (Extended Data Fig. 4d and Supplementary Table S4). Cytotoxic effectors like granzymes, perforin and Ifng were commonly upregulated, while the negative regulators of effector responses such as Tcf7, Slamf6, Pdcd4 and Cd5 were commonly downregulated^72, 73^ (Fig. 4d). Several other functional proteins including TCR/cytokine signaling (Cd247, Cd3e, Cd3d, Cd3g, Tcra, Cd28, IL12rb1/2, Il17ra, Il2ra/b, Ifnar1, Il21r, Jak3 and Stat3), stimulatory or inhibitory checkpoint molecules (Tnfrsf21, Ctla4, Icos, Tigit, Tnfrsf18, Havcr2, Lag3, Tnfrsf8 and Tnfrsf9), and transcription factors for T cell activation (Jun, Junb, Fosl2, Irf8 and Batf) increased in at least one condition (Fig. 4d and Supplementary Table 4). Of note, several negative regulators of anti-tumor responses were down-regulated in Cul5 KO cells including exhaustion-associated biomarkers (Nr4a2, Pdcd1 and Cd160), myeloid-derived suppressive cell promoting cytokine Csf2, and pro-apoptosis factor Bcl2l11. Although Tcf7 as a key transcription factor for memory cell formation reduced in Cul5 KO cells, Sell as another marker of memory precursor increased significantly upon TCR stimulation compared to NC cells^74^. Flow cytometry was performed to confirm the MS results on several critical markers of tumor immune responses including CD25 (Il2ra), CD5, CD137 (Tnfrsf9), ICOS, PD1 (Pdcd1), CTLA4, and CD62L (Sell) (Fig. 4e,f,g). In addition, after 16-hour stimulation, the number of live Cul5 KO cells increased, whereas that of NC cells decreased, compared to pre-stimulation (Fig. 4h). These results together are consistent with the overall enhanced persistence and cytotoxic activity of Cul5 KO T cells.

**Fig. 4:**
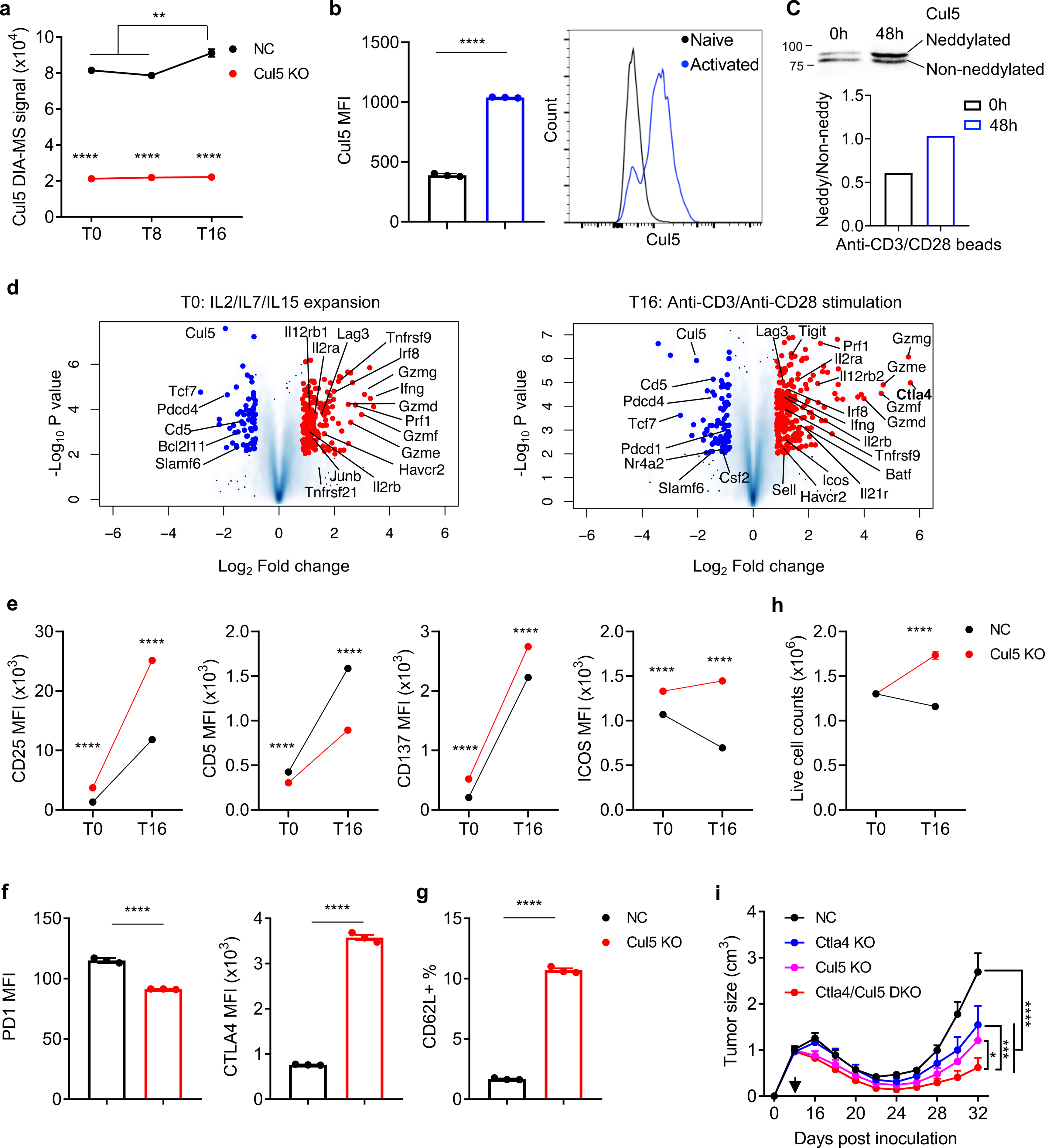
Cul5 KO causes proteomic alterations in primary CD8^+^ T cells. **a,** DIA-MS signals of the CUL5 protein in NC (Black) or Cul5 (Red) KO primary CD8^+^ T cells at the T0, T8 and T16 conditions determined by mass spectrometry (MS). Data are shown as mean ± SEM (**p < 0.01 and ****p < 0.0001 by multiple t-test, n=3). **b,** Flow cytometry analysis of Cul5 expression in primary CD8^+^ T cells before (Black) or after (Blue) 2-day activation by anti-CD3/CD28 beads. Data are shown as mean ± SEM (****p < 0.0001, by unpaired t-test, n=3). **c,** Western blot analysis of CUL5 expression in primary CD8^+^ T cells treated as in **b**. The ratio of neddylated to non-neddylated Cul5 (Bottom) was quantified. **d,** Volcano plot of differentially expressed proteins between Cul5 KO and non-targeting (NC) primary CD8^+^ T cells at T0 (Left) and T16 (Right) conditions as quantified by DIA-MS (n=3). **e,f,** Flow cytometry validation of immunological markers (CD25, CD5, CD137, ICOS, PD1, CTLA4 and CD62L) altered in Cul5 KO primary CD8^+^ T cells at T0 and/or T16 conditions. Data are shown as mean ± SEM (****p < 0.0001 by unpaired t-test, n=3). **h,** Live cell numbers of NC (Black) and Cul5 (Red) KO primary CD8^+^ T cells before and after 16-hour anti-CD3 plus anti-CD28 stimulation. Data are shown as mean ± SEM (****p < 0.0001 by unpaired t-test, n=3). **i,** Growth curve of tumors from E.G7-OVA cells inoculated s.c. into C57BL/6 mice. Data are shown as mean ± SEM. The back arrow indicates the time of sub-lethal irradiation followed by immediate adoptive transfer of Cas9/OT-I cells with NC (Black), Ctla4 (Blue), Cul5 (Pink) and Ctla4/Cul5 double (Red) KO. (*p<0.05, ***p<0.001 and ****p<0.0001 by two-way ANOVA; n=4-5).

We also noticed that several co-inhibitory checkpoint molecules increased along with the enhanced anti-tumor functions of Cul5 KO cells. These changes may reflect negative feedback regulation as the results of elevated T cell activation in the Cul5 KO cells. Among these molecules, the Ctla4 protein content showed the strongest increase in Cul5 KO cells (Fig. 4d,f). Therefore, we reasoned that knockout of Ctla4 in combination of Cul5 KO may further release the cytotoxic power of anti-tumor CD8^+^ T cells. To this end, in the same E.G7-OVA tumor model, we compared the anti-tumor effects of NC, Ctla4 KO, Cul5 KO and Ctla4/Cul5 double KO (DKO) OT-I T cells post adoptive transfer. Ctla4 KO alone showed similar ability in tumor control to Cul5 KO. However, as anticipated, Ctla4 and Cul5 DKO showed a superior anti-tumor ability compared to either Ctla4 KO or Cul5 KO alone (Fig. 4i and Extended Data Fig. 4e). This result suggests a therapeutic potential of ACT with a combinatory Cul5 and Ctla4 knockout.

### Cul5 interacts with Pcmtd2 and targets TCR/IL2 signaling

To identify Cul5 interacting proteins in CD8^+^ T cells, we overexpressed Cul5 with a C terminal HA-tag (Cul5-HA) in mouse primary CD8^+^ T cells by retroviral transduction. The cells were subjected to TCR stimulation for 12 hr, and Cul5-HA was immunoprecipitated by anti-HA, followed by DIA-MS analysis (co-IP-MS). Compared to the negative control samples (cells transduced with the empty vector), 65 proteins were enriched (p value <0.05 and fold change >1.5) in the anti-HA IP samples (Supplementary Table 5). Cul5 and the obligate Cul5 E3 ligase complex subunits including Rnf7, Neddy8, EloB and EloC were highly enriched (Fig. 5a). Among all of the known SOCS-box-containing proteins that have the potential to interact with Cul5 and were detectable in the total protein MS analysis (Fig. 5b), Pcmtd2^75^ is the only one enriched in the Cul5-HA IP samples (Fig. 5a). To confirm the importance of Pcmtd2 in TCR/cytokine signaling, we performed Pcmtd2 KO in mouse primary CD8^+^ T cells followed by cytokine-induced expansion/differentiation and TCR-induced re-stimulation. Pcmtd2 KO efficiency was confirmed by Western blotting analysis (Fig. 5b). Consistent to Cul5 KO, the expression of GZMB and IFNg was significantly higher in Pcmtd2 KO CD8^+^ T cells compared to NC (Fig. 5c), suggesting that Pcmtd2 functions as a Cul5 adaptor protein negatively regulating the differentiation and activation of effector CD8^+^ T cells. Of note, Asb6, which is a validated Cul5 binding protein^76^, was readily detected in CD8^+^ T cells (Extended Data Fig. 5a), but not enriched in the Cul5 co-IP samples, suggesting a possible selectivity of the Cul5/Eloc/Elob complex for Pcmtd2 in mouse CD8^+^ T cells. The lack of detection of Wsb1, Asb3, Socs3 or Socs1 in the Cul5-HA IP samples may be due to low abundance of these proteins or selectivity in CD8^+^ T cells (Extended Data Fig. 5a). Cish binds to Cul5 in CD4 T cells^41^ and was upregulated upon TCR stimulation in CD8^+^ T cells, but not enriched in Cul5-HA IP sample either, suggesting a possible differential utilization of substrate receptors between CD4^+^ and CD8^+^ T cells.

**Fig. 5:**
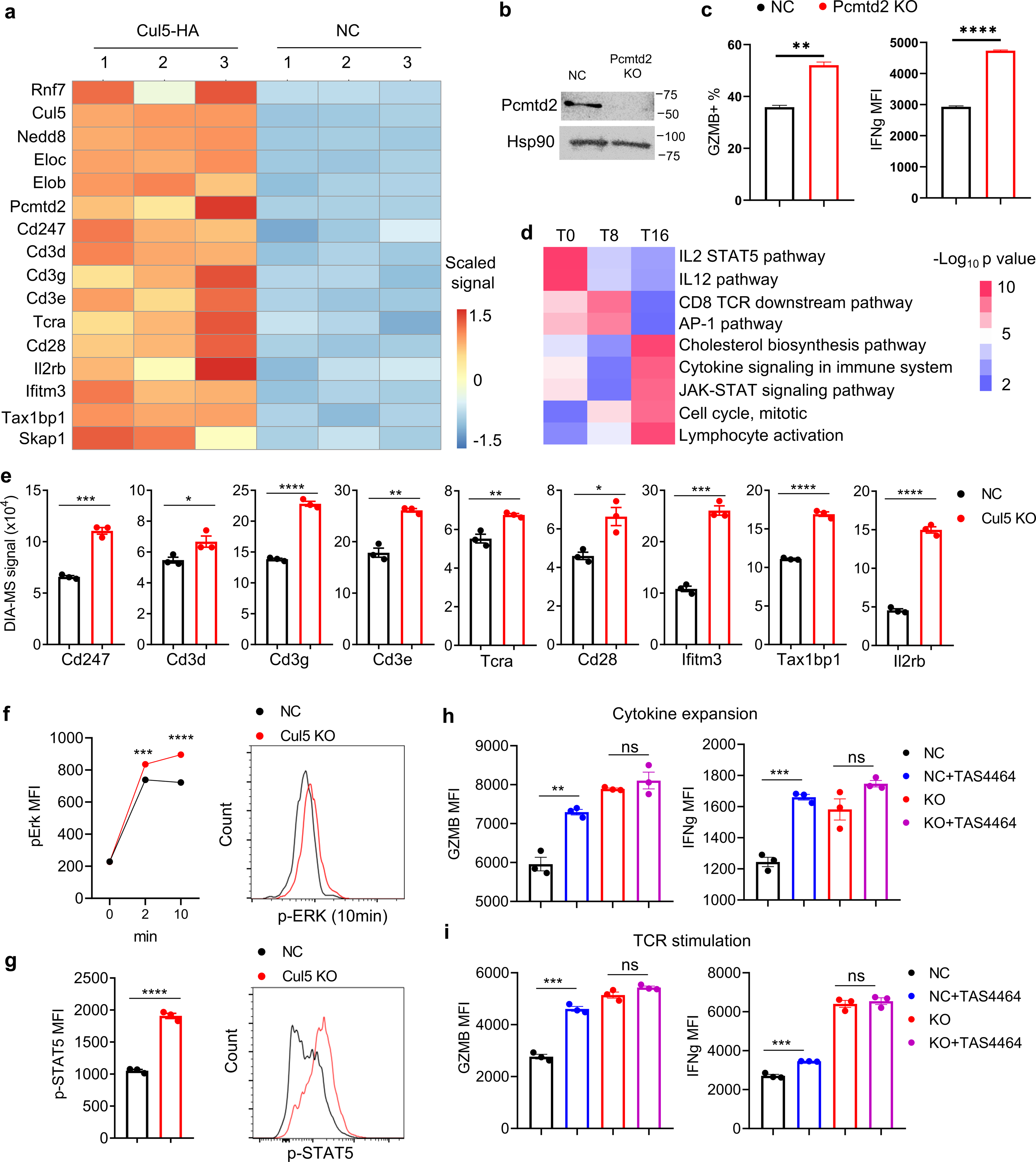
The Cul5 E3 complex targets TCR and IL2 signaling in CD8^+^ T cells. **a,** Heatmap of enriched proteins in TCR-activated mouse primary CD8^+^ T cells transduced with Cul5-HA overexpressing vector (Cul5-HA) vs empty control vector (NC) identified by anti-HA co-IP MS (Cut-off: p value <0.05 and fold change >1.5 in multiple t-test). **b,** Western blot analysis of Pcmtd2 expression in primary mouse CD8^+^ T cells with NC or Pcmtd2 KO. Hsp90 as an internal control. **c,** Flow cytometry analysis of GZMB (as positive % in cytokine expansion condition, Left) and IFNg (as MFI in 6-hour anti-CD3/CD28 stimulation condition, Right) expression of NC (Black) or Pcmtd2 KO (Red) primary mouse CD8^+^ T cells. Data are shown as mean + SEM (**p < 0.01 and ****p < 0.0001 by unpaired t-test, n=3). **d,** Heatmap of signaling pathway enrichment of Cul5 KO CD8^+^ T cells compared to NC cells based on differentially expressed proteins using DIA-MS quantification. **e,** DIA-MS signals of proteins identified by the total protein MS analysis of mouse primary Cul5 KO (Red) and NC (Black) CD8^+^ T cells post TCR stimulation. Data are shown as mean ± SEM (*p<0.05, **p < 0.01, ***p<0.001 and ****p < 0.0001 by unpaired t-test, n=3). **f,** Flow cytometry analysis of p-ERK1/2 expression in NC or Cul5 KO primary mouse CD8^+^ T cells after anti-CD3 plus anti-CD28 stimulation. Data are presented (Left) as mean ± SEM (***p<0.001 and ****p<0.0001 by unpaired t-test, n=3). **g,** Flow cytometry analysis of p-STAT5 expression in NC or Cul5 KO primary mouse CD8^+^ T cells after 16-hour anti-CD3 plus anti-CD28 stimulation supplied with 5ng/ml hIL2. Data are presented (Left) as mean ± SEM (****p<0.0001 by unpaired t-test, n=3). **h,i,** Flow cytometry analysis of GZMB and IFNg in NC (Black and Blue) or Cul5 KO (Red and Purple) primary CD8^+^ T cells in 16-hour **h,** cytokine culture (IL2/IL7/IL15) or **i,** anti-CD3 plus anti-CD28 stimulation with (Blue and Purple) or without (Black and Red) 250nM TAS4464. Data are shown as mean ± SEM (ns as not significant, **p < 0.01, ***p < 0.001, and ****p < 0.0001 by unpaired t-test, n=3).

Several other proteins related to T cell activation and responses were also significantly enriched in the Cul5-HA IP samples (Fig. 5a), including the TCR complex molecules (Cd3 subunits and Tcra), co-stimulator Cd28, IL2 receptor β subunit (Il2rb), Ifitm3 (supporting resident memory T cell survival)^77^, Tax1bp1 (important for cell-cycle and mTORC signaling)^78^, and Skap1 (important for LFA-1 adhesive activation)^79, 80^. Because these proteins were upregulated in Cul5 KO CD8^+^ cells (Fig. 5e), among which the surface expression of TCR complex and Il2rb (Extended Data Fig. 5b) were further confirmed by flow cytometry, they may be direct substrates of the Cul5 E3 ligase complex for ubiquitinoylation and subsequent degradation. Moreover, consistent with the importance of these molecules in T cell signaling, enrichment of a broad range of signaling pathways including the IL2/STAT5 signaling, AP1, CD8 TCR downstream signaling, IL12 signaling, cholesterol biosynthesis, TNF-alpha signaling via NF-kB, mTORC1 signaling, interferon gamma response, and cell cycle regulation pathways in Cul5 KO CD8^+^ T cells was revealed by pathway analysis of differentially expressed proteins based on the total protein MS data (Fig. 5d and Extended Data Fig. 5c). Increased Erk1/2 phosphorylation upon TCR stimulation (Fig. 5f) and STAT5 phosphorylation (Fig. 5g) in Cul5 KO CD8^+^ T cells further supports increased TCR downstream signaling and IL2/STAT5 signaling in Cul5 KO CD8^+^ T cells, which is consistent with the MS results.

Because Cul5 neddylation is required for the E3 ligase activity of the Cul5 complex, we tested a high-affinity neddylation inhibitor TAS4464^81^ on Cul5 KO and control (NC) CD8^+^ T cells. Although there are other cullin proteins expressed in CD8 T cells (Extended Data Fig. 5d) whose activity also depends on neddylation ^82^, TAS4464 increased GZMB and IFNg expression only in NC CD8^+^ T cells but not in Cul5 KO cells in both cytokine- (Fig. 5h) and TCR- (Fig. 5i) activation conditions, suggesting these two effector molecules are specifically regulated by Cul5, but not other cullin proteins in these cells. However, TAS4464 increased CD25 expression in both NC and Cul5 KO CD8^+^ T cells, suggesting other neddylated proteins may also negatively regulate CD25 in these cells (Extended Data Fig. 5e,f).

### CUL5 is a negative regulator of human CD8^+^ T cells

To evaluate the human relevance and translational potential of our findings, we studied CUL5 changes at protein level in primary human CD8^+^ T cells. The total CUL5 protein detected by both flow cytometry (Extended Data Fig. 6a) and western blot (Extended Data Fig. 6b) was significantly upregulated upon TCR-stimulation. CUL5 neddylation also increased (Extended Data Fig. 6b), consistent with the phenotype of mouse primary CD8^+^ T cells. We further performed CUL5 KO in primary human CD8^+^ T cells using the Cas9/sgRNA ribonucleoprotein electroporation approach. Three days post electroporation, CUL5 KO efficiency was evaluated by Western Blotting (Fig. 6a). Compared to the NC cells, CUL5 KO CD8^+^ T cells expressed significantly higher GZMB both before and after TCR-dependent activation (Fig. 6b). The percentage of IFNG^+^ cells and IFNG expression level are significantly higher in the CUL5 KO CD8^+^ T cells than the NC cells upon TCR-stimulation (Fig. 6c). There was no IFNG response or difference in CUL5 KO CD8^+^ T cells compared to the NC cells in the absence of TCR-stimulation (Extended Data Fig. 6c), suggesting CUL5 KO may not cause autoreactivity and tonic signaling in CD8^+^ T cells, thus minimizing the potential safety and accelerated exhaustion issues in clinical use. We further generated CD8^+^ chimeric antigen receptor (CAR)-T cells targeting CD19 by lentiviral transduction and sorted pure CAR-expressing cells by GFP reporter (Extended Data Fig. 6d). CUL5 KO CAR-T cells expressed significantly higher GZMB (Fig. 6d) and showed higher percentage of IFNG^+^ cells and IFNG expression level (Fig. 6e) after co-culturing with CD19-expressing NALM6 B cells, suggesting an increased CAR signaling activity in CUL5 KO CAR-T cells. Consistently, the CUL5 KO CAR-T cells showed significantly higher tumor cell killing ability of NALM6 B cells in culture than the NC cells (Fig. 6f). Moreover, as shown in murine CD8^+^ T cells, CUL5 KO also resulted in significantly higher CTLA4 expression in human CD8^+^ T cells than the NC cells upon TCR re-stimulation suggesting a potential combinatory CAR-T therapy with CTLA4 blockade (Fig. 6g). To further access the clinical relevance of CUL5 in cancer, we applied the Tumor Immune Dysfunction and Exclusion (TIDE) analysis to examine the association of CUL5 expression^83^ with T cell dysfunction and disease outcomes. In multiple cancer types, including ovarian cancer (Fig. 6h), leukemia (Fig. 6i), and melanoma (Fig. 6j), high cytotoxic T cell score is associated with an overall survival benefit only when CUL5 expression is low, while high CUL5 expression abolishes and even reverses the beneficial effects of infiltrating cytotoxic T cells. Taken together, our data suggest that CUL5 is expressed in primary human CD8^+^ T cells and plays important roles in CD8 cell activation and tumor killing functions in both TCR- and CAR-dependent ways.

**Fig. 6:**
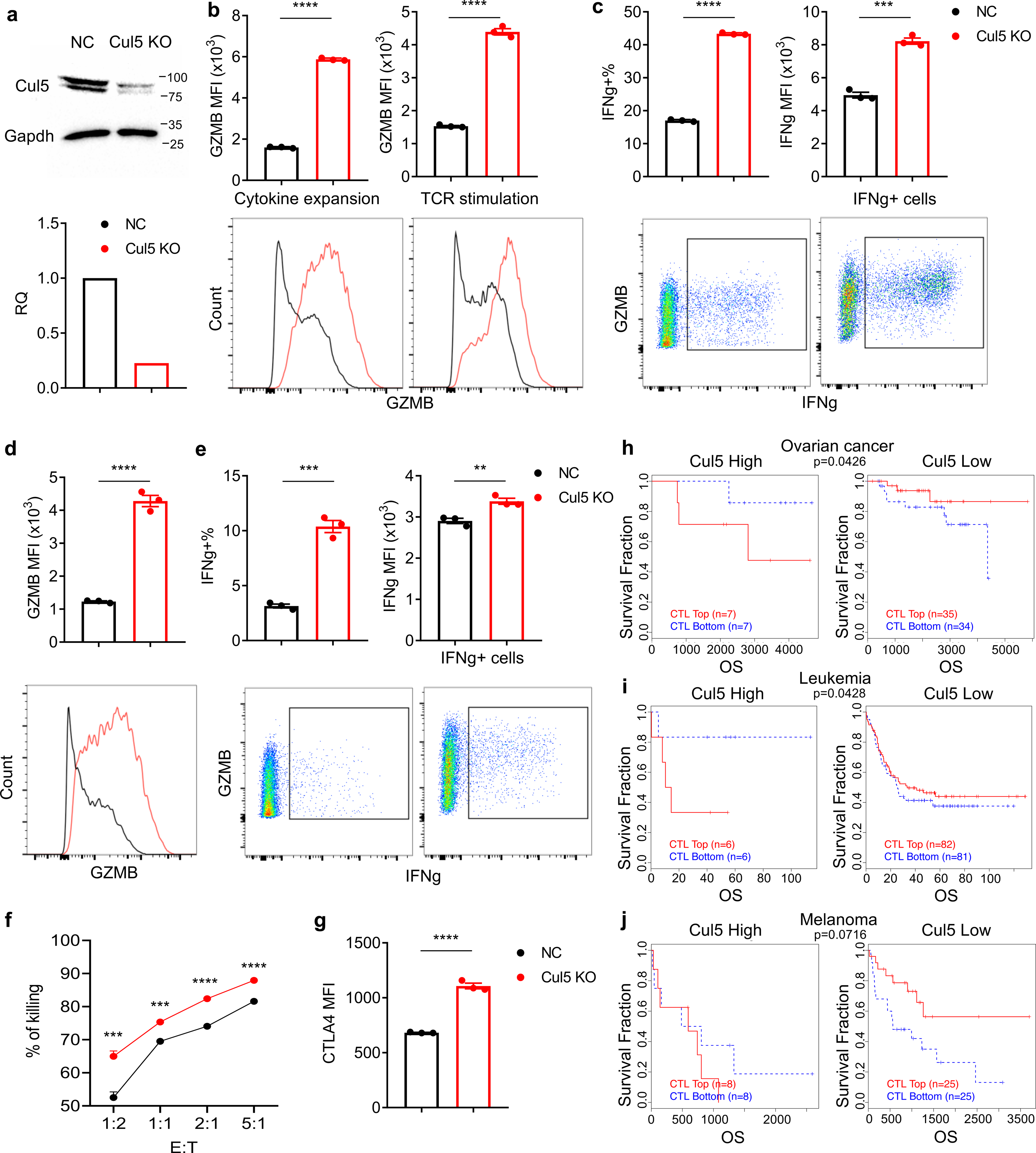
Characterization of CUL5 KO primary human CD8^+^ T cells. **a,** Western blot analysis of CUL5 expression in primary human CD8^+^ T cells with NC (Black) or CUL5 KO (Red). Quantification of CUL5 protein levels normalized against GAPDH is shown (Bottom). **b,** Flow cytometry analysis of GZMB expression in NC or CUL5 KO primary human CD8^+^ T cells before (Left) or after (Right) 6-hour anti-CD3 plus anti-CD28 stimulation. Data are shown in bar chart (Top) as mean ± SEM (****p<0.0001 by unpaired t-test, n=3). **c,** Flow cytometry analysis of IFNG^+^ cells and expression in NC (Black) or CUL5 KO (Red) primary human CD8^+^ T cells after 6-hour anti-CD3 plus anti-CD28 stimulation. Data are shown in bar chart (Top) as mean ± SEM (***p<0.001 and ****p<0.0001 by unpaired t-test, n=3). **d,e,** Flow cytometry analysis of **d,** GZMB expression as MFI and **e,** IFNG expression as positive % in total (Left) and MFI in positive population (Right) in NC (Black) or CUL5 KO (Red) primary human CD8^+^ T cells transduced with CAR-CD19 after 6-hour co-culture with NALM6 B cell line. Data are shown in bar chart (Top) as mean ± SEM and as representative histogram (**d**, Bottom) or scatter plot (**e,** Bottom) **p<0.01, ***p<0.001 **** and p<0.0001 by unpaired t-test, n=3. **f,** *In vitro* killing assay of NC (Black) or Cul5 KO (Red) CAR-CD19 primary human CD8^+^ T cells co-cultured with NALM6 B cell line overnight. E:T, T cell to NALM6 B cell ratios. The data presented as % of killing (mean ± SEM; ***p < 0.001 and ****p < 0.0001 by multiple t-test, n=3). **g,** Flow cytometry analysis of CTLA4 expression in NC or CUL5 KO primary human CD8^+^ T cells 2-day post anti-CD3 plus anti-CD28 stimulation. Data are shown as mean ± SEM (****p < 0.0001 by unpaired t-test, n=3). **h-j,** TIDE analyses of CUL5 expression correlation with T cell dysfunction signatures and survival benefits in patients with **h,** ovarian cancer, **i,** leukemia and **j,** melanoma. (Left panels: patients with high Cul5 expression; Right panels: patients with low Cul5 expression).

## Discussion

In this study, we used a step-wise CRISPR/Cas9 screens in combination of proteomic analyses to identify a Cul5-E3 ligase complex as an important negative feedback regulator of TCR/IL2 signaling and anti-tumor effector activity in CD8^+^ T cells. The Cul5 protein was significantly upregulated upon TCR stimulation, and its KO in CD8^+^ T cells significantly enhanced their effector function, TCR and cytokine signaling, stemness, and survival accompanied with improved tumor control ability *in vitro* and *in vivo* (Extended Data Fig. 7). To date, different screening studies of CD8^+^ T cells have highlighted various candidates based on diverse *in vitro* and/or *in vivo* models and different readout criteria such as proliferation^21, 23–25, 27^, effector molecule (degranulation or cytokine) expression^21, 22, 63^ or subpopulation^26, 84^. However, Cul5 was not identified as an enrichment hit in any of these studies, suggesting either enriched regulators in each screening are highly context-dependent or low-coverage in vivo screens tend to miss critical hits. Our step-wise screening strategy combining specific selection pressures with high confident coverages described in this study allowed identification of new hits including Cul5 that are important for signaling and anti-tumor activities of CD8^+^ T cells. Thus, this strategy provides a robust and cost-labor effective approach and can be readily adapted for other loss- or gain-of function screens with different selection pressures.

Cul5 E3 ubiquitin ligase-dependent protein degradation has been proposed as a mechanism by which the CIS/SOCS family proteins regulate T cell activation^39, 40^. However, our study unexpectedly identified Pcmtd2, rather than other CIS/SOCS family proteins, as the sole substrate receptor for Cul5 in mouse CD8^+^ T cells. Pcmtd2 has 79% amino acid sequence identity to its paralog Pcmtd1^85^. Both the Pcmtd molecules contain a Protein-L-Isoaspartate (D-aspartate) O-Methyltransferase domain and a SOCS-box domain. Pcmtd1 can bind to the canonical methyltransferase cofactor S-adenosylmethionine (AdoMet) and Cul5/EloB/EloC complex, but does not appear to have the L-isoaspartyl methyltransferase activity ^85^. Pcmtd2, which also can bind to Cul5 ^75^, but not Pcmtd1, was detected in our Cul5-HA co-IP MS analysis, suggesting Pcmtd2 may be the dominant isoform in CD8^+^ T cells. This may be due to low expression of Pcmtd1 as it was undetectable in our total protein MS analysis of CD8^+^ T cells. The function of these Pcmtd molecules is not known, even though Pcmtd1 was implicated in aging/stress-related protein repair^85^. The IP-MS results and those showing that Pcmtd2 KO in CD8^+^ T cells phenocopied Cul5 KO indicate that Pcmtd2 is the substrate receptor for Cul5 in CD8^+^ T cells and hence reveal the first known in vivo function for the Pcmtd proteins.

The Cul5-HA co-IP MS analysis also revealed an enrichment of the TCR/CD3 complex subunits, Cd28, Il2rb, Ifitm3 and Tax1bp1. These proteins also increased in Cul5 KO CD8^+^ T cells, suggesting that they may be direct target substrates of the Cul5 E3 ligase. TCR, CD28 co-stimulation and cytokines provide three signal traits that together induce optimal T cell activation. IL2 is critical for CD8^+^ T cell activation, proliferation and cytotoxic effector differentiation^48^ while IL15 is indispensable for the maintenance of memory CD8^+^ T cells^86^. Both cytokines signal through Il2rb. Ifitm3 increases upon T cell activation and maintains T cell survival and effector functions in the lung tissue with influenza infection^77^. It is also constitutively expressed in tissue-resident memory T cells, suggesting its role in the memory survival. Tax1bp1 acts through autophagy induction in activated T cells to support mTORC activation and subsequent T cell proliferation^78^. Upon TCR-induced CD8^+^ T cell activation, the expression of Cul5 and its neddylation significantly increased, rendering Cul5 as a negative feedback regulator for TCR and IL2/IL15 signaling. Consistent with this idea, Cul5 KO in CD8^+^ T cells resulted in an upregulation of TCR and cytokine/JAK/STAT signaling, which may subsequently increase the activity and expression of T cell-activation related TFs, biogenesis and metabolism enzymes, and cytotoxic effector molecules, leading to enhanced anti-tumor activity. Beyond the enhanced effector functions, elevated Jun^87^ and Batf^88^ and decreased Nr4a2 protein contents may explain increased exhaustion resistance^89, 90^. Although some memory/stemness associated markers such as Tcf7 and Slamf6 were reduced upon Cul5 KO, CD62L^+^ population in activated T cells associated with memory phenotype increased, consistent with increased persistence and IL7R expression of effector cells *in vivo* and elevated live cell number post re-stimulation *in vitro*. These together suggest that Cul5 KO may not compromise stem-like differentiation.

It is interesting that two well-characterized immune checkpoint molecules, PD-1 and CTLA4, changed in opposite directions in Cul5 KO CD8^+^ T cells. PD-1 expression decreased, whereas Ctla4 increased, in the Cul5 KO CD8^+^ T cells compared to the control cells. The reduction in PD-1 expression may reduce its immunosuppression and further facilitate activation of CD8^+^ T cells by Cul5 KO in tumors. The downregulation of PD-1 may be due to the increase of Satb1 expression in the Cul5 KO CD8^+^ T cells, as Satb1 is a direct suppressor of PD-1^91^ and negatively regulated by TGF-β^91, 92^. The marked increase in CTLA4 expression in activated Cul5 KO CD8^+^ T cells, on the other hand, should suppress Cul5 KO CD8^+^ T cells by B7-expressing cells in tumor. Indeed, CTLA4/Cul5 DKO in CD8^+^ T cells showed further improved tumor control *in vivo* compared to Cul5 KO alone. This combinatory therapeutic potential of transferred CD8^+^ T cells with Cul5 KO may apply to other Cul5 KO-induced co-stimulatory molecules (ICOS and 4-1BB) by administration of their agonists or to co-inhibitory molecules (Lag3, TIGIT and Tim3) by their blockade. More importantly, the characteristics of Cul5 KO were translational from mouse to human CD8^+^ T cells. Namely, Cul5 KO enhanced TCR- or CAR-dependent T cell activation and improved cytokine-induced effector T cell differentiation in human CD8^+^ T cells. In addition to ACT, it is also possible that Cul5 is a potential immunotherapeutic target for small molecule intervention, as small molecule inhibitors for E3 ligases such as neddylation inhibitors are being actively investigated ^93^. The observation that the neddylation inhibitor enhances CD8^+^ T cell activation and effector responses largely dependently of Cul5 supports these ideas and potential. Therefore, the Cul5-E3 ligase inhibitors may be tested as a single or combination with other ICIs, particularly anti-Ctla4 antibodies, giving high priority to the tumors in which Cul5 expression shows positive correlations with the clinical CTL dysfunction signature. Since neddylation inhibitors are being tested clinically, attentions should be given to assess immunological responses of the treatment.

## Methods

### Cell lines and mice

HT-2, HEK293T, EL4, E.G7-OVA and NALM6 cell lines were purchased from ATCC. YUMM1.7 cell line was kindly provided by M.B. and was reported previously. MC38 cell line was purchased from Kerafast (Boston, MA). All cell lines were regularly tested for mycoplasma and confirmed to be negative. OT-I TCR transgenic mice (OT-I mice) and Constitutive Cas9-expressing mice (Cas9 mice) both with C57BL/6 background were purchased from Jackson lab. Cas9/OT-I mice were generated by crossing Cas9 and OT-I mice with genotyping following Jackson Lab protocols. 7-8-week-old female C57BL/6N mice purchased from Envigo were used as E.G7-OVA tumor and adoptive T cell transfer recipients. Mice were housed in specific-pathogen-free conditions with all procedures approved by the Yale University Animal Care and Use Committee.

### Plasmids

The human Brie genome-wide CRISPR knockout pooled library in the pLentiCRISPRv2 one vector system (co-expressing spCas9 and sgRNA), with four sgRNAs per gene, was obtained from Addgene (Addgene # 73632, a gift from David Root and John Doench ^53^) and prepared in the Yale Cancer Center Functional Genomics core). pMSCV-U6sgRNA(BbsI)-PGKpuro2ABFP was a gift from Sarah Teichmann (Addgene plasmid # 102796 ; http://n2t.net/addgene:102796 ; RRID:Addgene_102796). PGKpuro2ABFP was replaced by EF1a-core promoter-mScarlet gblock (IDT) to generate pMSCV-guide-EF1a-mScarlet vector compatible with Bio-rad S3e sorting. For single cell perturb-seq, the original gRNA scaffold sequence was further replaced by a new scaffold sequence with 10x genomics compatible capture sequence 1 incorporated into the stem-loop to generate pMSCV-scguide-EF1a-mScarlet vector. Customized sub-library for bulk *in vivo* screening, with non-targeting control sgRNA sequences published previously and top ranked 6 sgRNA sequences per gene designed by CRISPick, was synthesized by Genscript and cloned into pMSCV-guide-EF1a-mScarlet. Customized sub-library for single cell *in vivo* screening, with 4 non-targeting control sgRNA sequences and top ranked 3 sgRNA sequences per gene selected from the above bulk sub-library, was synthesized by IDT and cloned into pMSCV-scguide-EF1a-mScarlet by NEB stable competent cells. The normal distribution and integrity of sub-libraries were confirmed by illumine next-generation-sequencing. Top two ranked murine Cd8a and Cul5 sgRNAs (Supplementary Table 6) were designed by CRISPick, synthesized by IDT and cloned into pMSCV-guide-EF1a-mScarlet or pMSCV-scguide-EF1a-mScarlet vector by NEB stable competent cells. pSLCAR-CD19-BBz was a gift from Scott McComb (Addgene plasmid # 135992 ; http://n2t.net/addgene:135992 ; RRID:Addgene_135992). Cul5-HA overexpression retroviral plasmid was derived from in house MIGR-IRES-GFP vector. Mouse Cul5 open-reading frame with HA tag at its C terminal was amplified by PCR. cDNA of TCR-activated mouse primary CD8^+^ T cells was used as PCR template. Primers were designed to add XhoI site at 5’ and HA sequence plus EcoRI site at 3’ of the PCR product (Supplementary Table 6). Backbone vector and PCR product were double digested by XhoI and EcoRI, and ligated by T4 ligation to get MIGR-Cul5-HA-IRES-GFP plasmid.

### Lentiviral and retroviral production

Lentiviruses with genome-wide murine CRISPR library or CAR-CD19 were produced in HEK293T cells transfected with library vector, pMD2.G (addgene, #12259) and psPAX2 (addgene, #12260). Retroviruses with sub-libraries or individual non-targeting or target gene sgRNAs were produced in HEK293T cells transfected with sublibrary/guide vector and pCL-Eco (addgene, #12371). Viral soups from 24-hour and 48-hour post-transfection were harvested, combined, filtered, aliquoted and saved in -80 °C freezer until use. The titers of lentiviruses and retroviruses of each batch were determined by the transduction of HEK293T and NIH3T3 cells respectively.

### *In vitro* genome-wide CRISPR KO screening

HT-2 cells were transduced with the lentiviral CRISPR/Cas9 library at MOI of 0.3 with over 200x coverage per sgRNA. About 25% transduction efficiency was confirmed by three-day puromycin selection to make sure one type of sgRNA per cell as the majority (Extended Data Fig. 1a). After 3 days of selection with 0.6µg/ml puromycin, 2×10^7^ transduced HT-2 cells were saved as input, 2×10^7^ were cultured for 21 days either in 100ng/ml IL2 medium (IL2) or in IL2 medium containing 120pg/ml TGF-β1 (IL2+TGF). For each condition, genomic DNA from 2×10^7^ cells was extracted by DNeasy blood & tissue kit (Qiagen, Cat.69504) and sgRNA cassettes were PCR amplified for illumina NGS ^53^.

### *In vivo* bulk CRISPR KO screening

3×10^6^ EG.7-OVA cells were s.c. inoculated into 7-8 weeks old C56BL/6N female mice. Tumor sizes were monitored every three days by caliper with volume calculation as v=d^2^xD/2. When tumors reached 0.5 cm^3^, untouched CD8^+^ T cells were isolated from the spleens of Cas9/OT-I mice by CD8^+^ T cell isolation kit (MACS, Cat.130-104-075), and immediately stimulated by anti-mouse CD3/28 beads (Thermo Scientific, Cat.11456D) at 1:1 ratio for 24 hours, followed by retroviral sub-library transduction at MOI of 0.2. Transduced CD8^+^ T cells were further expanded in IL2/7/15 medium for 3 days to ensure genome-editing completed. GFP/mScarlet double positive cells were then sorted on a Bio-Rad S3e sorter. 2×10^6^ cells were saved as input and 2×10^6^ sorted T cells per mouse were i.v. transferred into a total of four sub-lethally irradiated (4Gy) EG.7-OVA tumor-bearing mice. 12 days post T cell transfer, GFP^+^ T cells were isolated and enriched from tumors, spleens and tumor-draining lymph nodes of recipient mice by sorting. Genomic DNA from sorted cells were extracted by Qiagen DNeasy blood & tissue kit. sgRNA cassettes were PCR amplified for illumina NGS.

### PCR of sgRNA cassettes for NGS

For all PCR products, a stagger P5-read1 forward primer mixture was used to increase NGS reading diversity, and different index-included P7-read2 reverse primers were used for PCR of individual replicate samples before pooled NGS. Primer sequences are shown in Table S6. PCR reaction was set up according to NGS protocol of NEBNext Ultra^TM^ II Q5 Master Mix (NEB, Cat.M0544S). PCR products were purified by AMPure XP beads (Beckman Coulter, Cat. A63880) at 1:1 volume ratio. Purified PCR products were quantified by TapeStation (D1000 ScreenTape assay) before loading for NGS.

### *In vivo* single cell CRISPR KO screening

Tumor inoculation and sub-library transduced T cell transfer followed the same procedures of *in vivo* bulk CRISPR KO screening. 7 days post T cell transfer, live transferred tumor-infiltrating CD8^+^ T cells as CD8a^+^GFP^+^ T cells were sorted and washed once in cold 1x PBS containing 0.04% BSA prior to resuspending in cold 1x PBS containing 0.04% BSA at 1×10^6^ cells/ml concentration. Cells were then processed by Yale Center for Genome Analysis to generate single-cell RNA library and sgRNA library separately according to the instruction of 10x genomics 3’ V2 single cell RNA sequencing kit with sgRNA feature. Libraries were sequenced by illumina NGS.

### *In vivo* gene KO validation

1×10^6^ sorted NC or Cul5 sgRNAs (Supplementary Table 6) transduced Cas9/OT-I T cells per mouse were i.v. transferred into sub-lethally irradiated (4Gy) EG.7-OVA tumor-bearing mice in total of 4-6 mice per group when average tumor sizes reached around 1 cm^3^. Tumor sizes were monitored every two days post T cell transfer. Ctla4 KO was achieved by nucleofection of ribonucleoprotein (RNP) complex containing Cas9 protein (IDT, Cat.1081058) and Ctla4 sgRNA (Supplementary Table 6) into naïve CD8^+^ T cells before activation. At the endpoint, tumors and TDLNs were isolated to make single cell suspension for immediate flow cytometry staining or *in vitro* re-stimulation by PMA/ionomycin prior to flow cytometry staining.

### Tissue processes and transferred CD8^+^ T cell isolation

EG.7-OVA tumors were resected and minced into 1-2mm pieces, followed by 20 minutes digestion at 37 °C in HBSS buffer containing Mg^2+^, Ca^2+^, HEPES, 2% FBS, collagenase, DNase I with constant shaking at 60 rpm speed. Digestion was stopped by 10mM ETDA PBS solution and pass through 70µm strainer. Remaining tumor pieces were further smashed with 3ml syringe plunger and washed through the strainer. For CD8^+^ T cell sorting, tumor single cell suspension in 1xPBS was layered on top of Ficol-plus and centrifuged at 1000g x 20minutes without braking at room temperature. Buffy coat in which live CD8 T cells were enriched was collected and washed in 1xPBS. Spleens and TDLNs were directly smashed with 3ml syringe plunger through 70µm strainers. Red blood cell lysis buffer was further applied for smashed splenocytes to remove red blood cells. Single cell suspensions from different tissues were stained with PE-anti-mouse CD8a. GFP^+^ CD8a^+^ cells were sorted by Bio-Rad S3e sorter.

### *In vitro* CD8^+^ T cell stimulation and cancer cell killing assay

Following retroviral transduction of MACS purified Cas9/OT-I T cells as described above, GFP^+^mScarlet^+^ cells were sorted and expanded in culture medium containing 5ng/ml hIL2 (R&D, Cat.202-IL-010), 2.5ng/ml mIL7 (Peprotech, Cat.217-17) and 25ng/ml mIL15 (Peprotech, Cat.500-P173) for 3-5 days. For flow cytometry detection of phosphorylated proteins, activation and effector markers, T cells at 1×10^6^/ml were kept in culture withdraw of cytokines for 8 hours before stimulated with 1µg/ml plate-coated anti-mouse CD3e (Biolegend, Cat.100340) and 1µg/ml soluble anti-mouse CD28 (Biolegend, Cat.102116) for different time points with or without addition of transporter block (Thermo Scientific, Cat.00-4980-03). For some experiments, 250nM TAS4464 (Selleckchem, Cat.S8849) was added for the inhibition of neddylation. Supernatants at different time points without the addition of transporter block were collected and kept in -80 °C freezer for cytokine ELISA detection later. For cancer cell killing assay, EL4 cells pre-stained with CFSE (Thermo Scientific, Cat.C34554) and EG.7-OVA cells pre-stained with CellTrace violet (Thermo Scientific, Cat.C34557) according to instruction were 1:1 mixed and seeded in U-bottom 96-well plate with 1×10^5^ total cells per well. T cells with different effector to target ratio were added and kept in culture overnight. Counting beads were added to calculate the absolute number of live CFSE^+^ EL4 cells and violet dye^+^ EG.7-OVA cells by flow cytometry. In some experiments, T cells and EL4/EG.7-OVA cells were co-cultured for 6 hours with the addition of transporter block to detect antigen-specific cytotoxic T cell activation by flow cytometry.

### Pcmtd2 KO in primary mouse CD8^+^ T cells

MACS-purified CD8^+^ T cells from spleens of WT mice were cultured in 5ng/ml mIL7 overnight. Then Pcmtd2 knockout of CD8^+^ T cells was achieved by CRISPR-Cas9. In brief, pre-designed crRNAs (IDT) for mouse Pcmtd2 (Supplementary Table 6) or non-targeting control was annealed with tracrRNA (IDT, Cat.1072532) to form guide RNA, which then was mixed with Cas9 protein to generate ribonucleoprotein (RNP) complex. 1×10^6^ CD8^+^ T cells were re-suspended in P3 buffer (Lonza) containing RNP complex and enhancer DNA (IDT, Cat.1075915) and under electroporation with CM-137 program of 4D-nucleofactor X Unit (Lonza). Electroporated T cells recovered overnight in IL7 medium were then activated by anti-mouse CD3/CD28 beads (at 1:1 ratio for two days and expanded in culture medium containing 5ng/ml hIL2, 2.5ng/ml mIL7 and 25ng/ml mIL15 for 2 days before experiments.

### RT-qPCR

RNA was extracted from cells by RNeasy plus mini kit (Qiagen, Cat.74034) and quantified by nanodrop. cDNA was then synthesized by iScript Reverse Transcription Supermix (Bio-Rad, Cat.1708841) for RT-qPCR from Bio-Rad. qPCR reaction was set up by using SsoAdvanced Universal SYBR Green Supermix (Bio-Rad, Cat.1725270) and run on CFX96 Real-Time PCR Detection System (Bio-Rad). Primers of individual genes were selected from PrimerBank as shown in Table S6 and synthesized by IDT. The specificity of qPCR reaction was confirmed by melting curve. Relative quantification of individual genes was normalized to Actb.

### ELISA

Supernatants of *in vitro* stimulated CD8^+^ T cells without secretion blocking were harvested at 16-hour post stimulation, and stored in the -80 °C freezer before test. Mouse IFNg ELISA (Thermo Scientific, Cat.KMC4021) were performed according to the instruction. In brief, capture antibodies were coated in 96-well plate overnight in 4 °C. Cytokine standards and sample supernatants were then added, followed by the addition of detection antibodies and secondary antibody-HRP conjugation. After extensive washes between each step, TMB was added for 20 minutes at room temperature in dark. After stopping the reaction, absorbance at 450nm was detected. IFNg concentrations were then calculated based on the standard curves.

### Flow cytometry

Cells were washed once in 1x cold PBS and resuspended in 50µl 1xPBS containing anti-CD16/32 (BD, Cat.553142) and fixable aqua live/dead dye (Thermo Scientific, Cat.L34957) for 10 minutes on ice. Without wash, 50µl surface antibody/tetramer mixture in 1xPBS was added for 15 minutes on ice, followed by 1x cold PBS wash twice. For intracellular cytokine, CTLA4 Cul5 and pERK1/2 staining, cells were fixed in 4% PFA for 20 minutes at room temperature and washed in 1x permeable buffer (Thermo Scientific, Cat.00-833356). 50µl intracellular antibody mixture in 1x permeable buffer was added for 30 minutes at room temperature, followed by 1x permeable buffer twice. 50µl anti-rabbit-PE or -Alexa Fluor 350 secondary antibody in 1x permeable buffer was added for 30 minutes at room temperature, followed by 1x permeable buffer twice. For intracellular pSTAT5 staining, cells were fixed in 4% PFA for 20 minutes at room temperature and washed in 1x PBS, followed by permeabilization in pre-colded methanol for 20 minutes on ice. After 1xPBS wash, 50µl anti-pSTAT5-APC in 1xPBS was added for 30 minutes at room temperature, followed by 1xPBS wash twice. After final wash, cells were resuspended in 1x PBS and detected by BD LSR II. Antibodies used for flow cytometry are as follows: anti-mouse: Biolegend: CD8a-PE-Cy7 (Cat.100721), CD122-APC (Cat.105911), CD127-APC-Cy7 (Cat.135039), GZMB-APC (Cat.372203), IFNg-APC-Cy7 (Cat.505849), TNF-PE-Cy7 (Cat.506323), IL2-PB (Cat.503820), CD25-APC-Cy7 (Cat.101917), CD5-PB (Cat.100641), ICOS-APC (Cat.107711), PD1-PE-Cy7 (Cat.135215), CTLA4-APC (Cat.106309), CD62L-PE-Cy7 (Cat.104417), Vb5-PB (Cat.139515), Va2-PE (Cat.127807); Thermo Scientific: CD137-PB (Cat.48-1371-82); BD: CD107a-APC (Cat.560646), pSTAT5-Alexa 647 (Cat.562076), anti-human (Biolegend): GZMB-APC (Cat.372203), IFNg-PE (Cat.502508), CTLA4-APC (Cat.369611), anti-rabbit: IgG-PE (Thermo Scientific, Cat. P-2771MP), IgG-Alexa Fluor 350 (Thermo Scientific, Cat.A-11069), anti-pERK1/2 (Cell Signaling Technology, Cat. 9101), Rabbit polyclonal IgG anti-Cul5 (Thermo Scientific, Cat.A302-173A); SIINFEKL-H-2K(b) tetramer-BV421 (NIH Tetramer Core Facility, Cat.53995)

### Proteomics analysis and co-IP for Mass Spectrometry

For proteomics measurements, 1×10^6^ cytokine expanded CD8^+^ T cells with or without Cul5 KO were either immediately harvested as T0 or continuously cultured in medium without cytokines for 8 hours. Then cytokine deprived CD8^+^ T cells were harvested either immediately as T8 or after 16 hours stimulation with 1µg/ml plate-coated anti-CD3 and 1µg/ml soluble anti-CD28 as T16. Harvested cells were washed in cold PBS for 3 times, snap frozen in liquid nitrogen and stored in -80 °C freezer. Cell pellets were lysed by the lysis buffer 10 M urea containing the cOmplete™ protease inhibitor cocktail (Roche, #11697498001). The cell tubes were ultrasonicated at 4 ℃ for two cycle (1 min per cycle) using a VialTweeter device (Hielscher-Ultrasound Technology)^94, 95^, and then centrifuged at 20,000 x g for 1 h to remove the insoluble material. For the supernatant protein mixture, the reduction and alkylation were conducted with 10 mM Dithiothreitol (DTT) for 1 h at 56 °C and then 20 mM iodoacetamide (IAA) in dark for 45 min at room temperature. The samples were diluted by 100 mM NH4HCO3 and digested with trypsin (Promega) at ratio of 1:20 (w/w) overnight at 37 °C. The digested peptides purification was performed on C18 column (MarocoSpin Columns, NEST Group INC) and 1 µg of the purified peptides was injected for mass spectrometry analysis.

For co-IP in triplicate, 1×10^7^ CD8^+^ T cells per replicate transduced with MIGR-IRES-GFP control vector or MIGR-Cul5-HA-IRES-GFP Cul5-HA overexpression vector were stimulated in 1ug/ml anti-CD3 coated plate plus 1ug/ml soluble anti-CD28 for 12 hours. 10µM MG-132 was added for the last 8 hours before harvesting cells. Harvested cells were washed in cold PBS for 3 times and immediately resuspended in IP lysis buffer (Thermo Scientific, Cat.87787) with proteinase/phosphatase inhibitor cocktail on ice for 10 minutes. After a high-speed centrifuge, supernatant was transferred into a new tube, followed by overnight incubation with anti-HA antibody (Biolegend, Cat.901515). Protein G dynabeads (Thermo Scientific, Cat.10003D) were then added with rotation at room temperature for 1 hour. Dynabeads containing bound protein complex were sequentially washed with cold IP lysis buffer without proteinase/phosphatase inhibitor cocktail 4x, and wash buffer without detergent 2x on magnetic stand. Protein complex was eluted twice by elution buffer containing 0.5M ammonium hydroxide and 0.5mM EDTA. Combined elutes were snap frozen in liquid nitrogen and stored in -80 °C freezer. The elution was dried with SpeedVac, and then resolved with 50 μL 6 M urea for reduction and alkylation. Other proteomic sample preparation steps were identical with the above cell sample protocol.

The samples were measured by data-independent acquisition (DIA) mass spectrometry method as described previously^96–98^. The Orbitrap Fusion Tribrid mass spectrometer (Thermo Scientific) instrument coupled to a nanoelectrospray ion source (NanoFlex, Thermo Scientific) and EASY-nLC 1200 systems (Thermo Scientific, San Jose, CA). A 120-min gradient was used for the data acquisition at the flow rate at 300 nL/min with the temperature controlled at 60 °C using a column oven (PRSO-V1, Sonation GmbH, Biberach, Germany). Each DIA-MS cycle consisted of one MS1 scan and 33 MS2 scans of variable isolated windows with 1 m/z overlapping between windows. The MS1 scan range was 350 – 1650 m/z and the MS1 resolution was 120,000 at m/z 200. The MS1 full scan AGC target value was set to be 2E6 and the maximum injection time was 100 ms. The MS2 resolution was set to 30,000 at m/z 200 with the MS2 scan range 200 – 1800 m/z and the normalized HCD collision energy was 28%. The MS2 AGC was set to be 1.5E6 and the maximum injection time was 50 ms. The default peptide charge state was set to 2. Both MS1 and MS2 spectra were recorded in profile mode. DIA-MS data analysis was performed using Spectronaut v15^99–101^ with “directDIA” by searching against the mouse SwissProt protein database. The oxidation at methionine was set as variable modification, whereas carbamidomethylation at cysteine was set as fixed modification. Both peptide and protein FDR cutoffs (Qvalue) were controlled below 1% by Spectronaut and the resulting quantitative data matrix were exported from Spectronaut. The sample-wise normalization was disabled for IP-MS experiments. All the other settings in Spectronaut were kept as Default.

### Human T cell culture and manipulation

CD8^+^ T cells were purified from human PBMCs by untouched human CD8^+^ T cell isolation kit (MACS, Cat.130-094-156) and cultured at 1×10^6^/ml in RPMI1640 medium containing 10% FBS, 5ng/ml hIL7 (R&D, Cat.207-IL-005) and 50ng/ml hIL15 (R&D, Cat.247-IL-005) overnight. Then Cul5 knockout of CD8^+^ T cells was achieved by CRISPR-Cas9. In brief, pre-designed crRNA (IDT) for human CUL5 (Supplementary Table 6) or non-targeting control was annealed with tracrRNA (IDT, Cat.1072532) to form guide RNA, which then was mixed with Cas9 protein to generate ribonucleoprotein (RNP) complex. 2×10^6^ CD8^+^ T cells were re-suspended in P3 buffer (Lonza) containing RNP complex and enhancer DNA (IDT, Cat.1075915) and under electroporation with EO-115 program of 4D-nucleofactor X Unit (Lonza). Electroporated T cells recovered overnight in IL7 and Il15 medium were then activated by anti-human CD3/CD28 beads (Thermo Scientific, Cat.11131D) at 1:1 ratio for two days. For CAR-T generation, one day post beads activation, T cells were transduced with lentiviruses containing CAR-CD19 by spinfection supplied with 8µg/ml polybrene and 5ng/ml hIL2. Activation beads were magnetically removed two days later and T cells at 5×10^5^/ml concentration were expanded in culture medium containing 5ng/ml hIL2, 5ng/ml hIL7 and 50ng/ml hIL15. Fresh medium with cytokines was replaced every two days to keep T cell concentration lower than 2×10^6^/ml.

### Western Blotting

WT mouse naïve CD8^+^ T cells were activation by anti-CD3/CD28 beads at 1:1 ratio for 0 or 48 hours. Human CD8^+^ T cells were activated by anti-CD3/CD28 beads at 1:1 ratio for two days and NC or CUL5 KO ones were further expanded in 5ng/ml hIL2, 5ng/ml hIL7 and 50ng/ml hIL15. Cells were then pelleted and washed twice with cold 1xPBS, followed by direct lysis in 2x SDS loading buffer (Thermo scientific, Cat.NP0007). Rabbit polyclonal IgG anti-Cul5 (Thermo Scientific, Cat.A302-173A) was used to detect mouse or human Cul5 expression. Rabbit polyclonal IgG anti-Pcmtd2 (Thermo Scientific, Cat. PA5-69557) was used to detect mouse Pcmtd2 expression. Mouse monoclonal IgG anti-GAPDH (Proteintech, Cat.60004-1-Ig), rabbit monoclonal IgG anti-Hsp90 (Proteintech, Cat. 13171-1-AP) and rabbit monoclonal IgG anti-Histone H3 (Cell Signaling Technology, Cat. 4499) were used as internal controls.

### Human CD8^+^ T cell activation *in vitro*

Cytokine expanded CD8^+^ T cells were stimulated in 1µg/ml plate-coated anti-CD3 and 1µg/ml soluble anti-CD28. CAR-T cells were stimulated with NALM B cells (ATCC, Cat.CRL-3273) at 1:2 ratio. Stimulated CD8^+^ T cells or CAR-T cells were cultured for 6 hours with the addition of transporter inhibitor prior to flow cytometry staining.

### Human CAR-T cell *in vitro* killing

T cells expressing CAR-CD19 were sorted based on GFP reporter. Sorted CAR-T cells were co-cultured with CellTrace violet stained NALM B cells at different effector to target (E:T) ratio overnight. Violet^+^ live NALM B cells were detected by flow cytometry with counting beads added to calculate absolute number. Killing % =(live B cells in B cell culture alone – live B cells in co-culture)/live B cells in B cell culture alone.

### Bioinformatics analysis

#### Bulk CRISPR KO screening analysis

Initial quality check was performed using the FastQC program and sequence adapters were trimmed using the Cutadapt tool. Genome-scale and enriched sub-library scale CRISPR/Cas9 KO screening performed using MAGeCK version 0.5.9.2^102^. MAGeCK uses a maximum likelihood based estimation (MLE) to measure Z-scores for each gene from the log2 fold changes of each sgRNAs in a robust approach. We rank our sgRNAs/genes based on robust ranking aggregation (RRA) and p-values. For bulk *in vivo* screening, four replicates in each group (Tumor, Spleen and Tumor-draining lymph node and Input) expression were averaged and positive RRA scores were used for the identification of top-ranked genes.

#### In vivo single-cell CRISPR KO screening analysis

Seurat 4.0 was used to process single-cell sequencing data. In the quality control (QC) analysis, poor quality cells with nFeature_RNA < 600, nCount_RNA < 1200, log10(Gene Per UMI) < 0.8 and Mitochondrial gene percentage over 10% were excluded. Genes with zero count numbers were removed. In addition, TCR genes were removed to prevent clustering bias caused by the contribution of variable V(D)J transcripts in major variable components. Cell cycle markers from Tirosh et al, 2015^103^ is loaded with Seurat and cell-cycle scores for G2M, S or G1 phase were quantified and assigned to each cell as the metadata. In addition, the sgRNA data was added into corresponding Seurat objects as the metadata. The cells with more than 1 sgRNA or without sgRNA were excluded. The feature-barcode matrix was first normalized and scaled with default settings in Seurat. Then the 3000 top variable genes were identified, which served as the input to principal component analysis (PCA) for dimensionality reduction. We applied Louvain algorithm for clustering and retained the 20 leading principal components as an input for further visualization. The UMAP embedding was used to visualize the single cells on a two-dimensional space with a perplexity of 100. Cell clusters were annotated with hall marker genes. Differentially expressed gene (DEG) analysis among different cell groups were based on the non-parametric Wilcoxon rank sum test with logfc.threshold=0.25 and min.pct =0.1. Functional analyses of sgRNA perturbations were evaluated by DEG analysis of sgRNA specific groups. Enrichr^104, 105^ was then used to perform signaling pathway enrichment analysis with The Molecular Signatures Pathway Database (MsigDB_Hallmark_2020) to discover enriched biological pathways based on the list of identified DEGs.

#### Proteomics data statistical analysis

The two-sided Student’s t-test was used to find the differentially abundant proteins, Protein groups with p < 0.01 and a fold-change > 1.8 were reported as significant in total protein MS while p<0.05 and a fold-change >1.5 in co-IP MS. The R software was used for the data virtualization with the packages ggplot2 (boxplot), scatterplot (volcano plots), pheatmap (correlation), corrplot (correlation), factoextra (PCA). The GO enrichment analysis was performed by Metascape^106^ with default Express Analysis.

All scripts for bioinformatics analysis will be available upon request.

## Data availability

The single-cell sequencing data has been deposited to GEO. The accession number is GSE213921. (To review the dataset please go to https://nam12.safelinks.protection.outlook.com/?url=https%3A%2F%2Fwww.ncbi.nlm.nih.gov%2Fgeo%2Fquery%2Facc.cgi%3Facc%3DGSE213921&data=05%7C01%7Cjingjing.ren%40yale.edu%7C138480356bee4bcf1dd908da9c98109b%7Cdd8cbebb21394df8b4114e3e87abeb5c%7C0%7C0%7C637994474170177851%7CUnknown%7CTWFpbGZsb3d8eyJWIjoiMC4wLjAwMDAiLCJQIjoiV2luMzIiLCJBTiI6Ik1haWwiLCJXVCI6Mn0%3D%7C3000%7C%7C%7C&sdata=s9wZzFbvvonyeGqr%2FzGP4J7MsuRpU6ReHqnUqd8a1%2BE%3D&reserved=0, with password: wfilwimmtbirpyl) All the mass spectrometry datasets and processed results have been deposited to the ProteomeXchange Consortium via the PRIDE partner repository with the dataset identifier PXD036793 (To review the dataset please go to https://www.ebi.ac.uk/pride/login, and use the following login details: Username, Password: EVCVbBJi).

---

**Extended Data Fig. 1.**
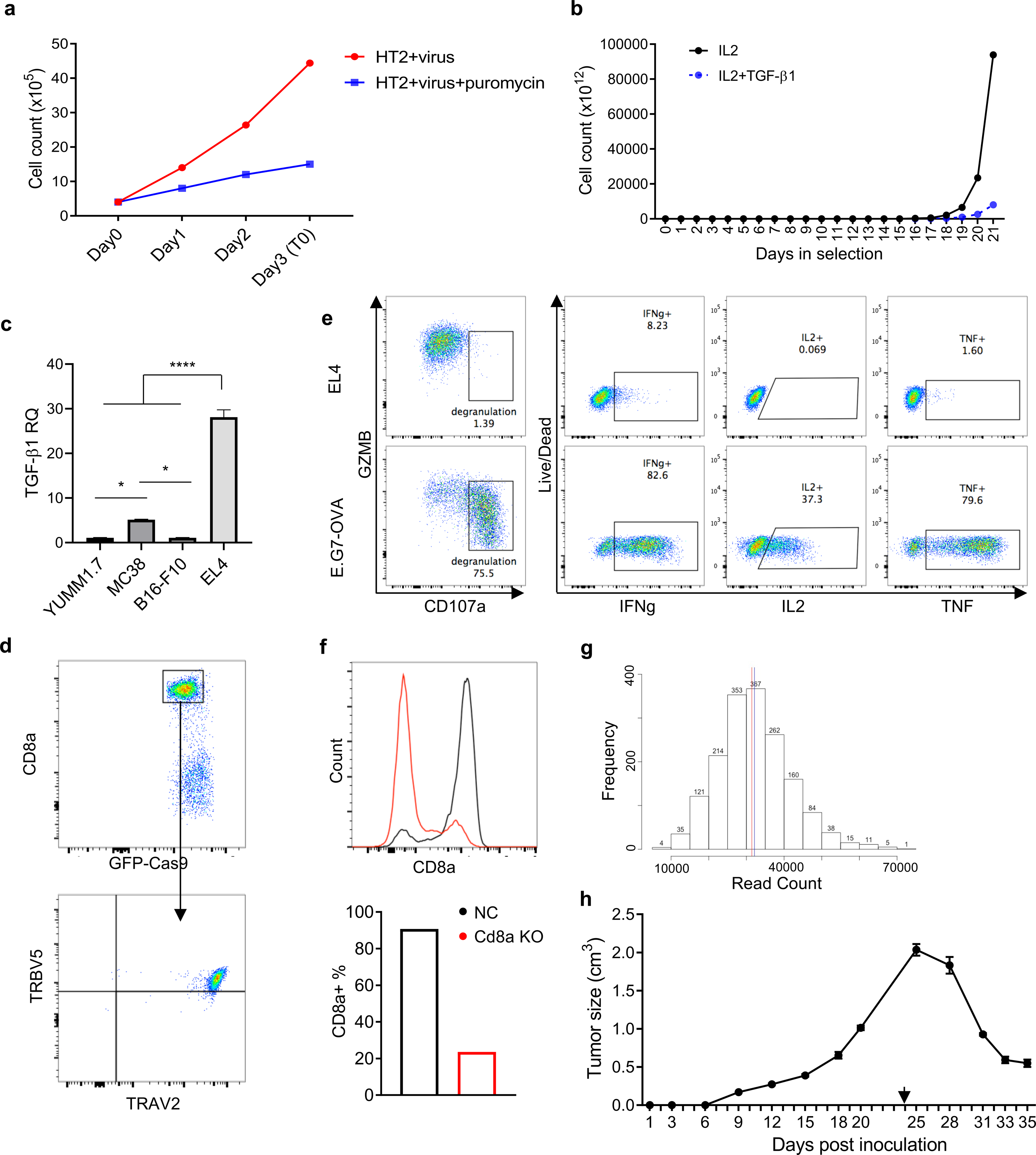
Bulk *in vitro* and *in vivo* CRISPR KO screens identify genes enhancing anti-tumor activity of CD8^+^ T cells. **a,** *In vitro* growth curve of HT-2 cells with (Blue) or without (Red) puromycin selection post library lentiviral transduction. **b,** *In vitro* growth curve of puromycin-selected HT-2 cells transduced with genome-wide CRISPR KO library in IL2 (Black) or IL2 plus TGF-β1 (Blue) culture condition. **c,** qPCR of Tgf-β1 mRNA level in different tumor cell lines. Data are shown as mean + SEM. *p < 0.05 and ****p < 0.0001 by one way ANOVA, n=3 **d,** Flow cytometry analysis of Cas9/OT-I T cells regarding GFP, TRBV5 and TRAV2 expressions. **e,** Flow cytometry analysis of the expression of activation markers (GZMB, CD107a, IFNg, IL2 and TNF) in Cas9/OT-I T cells post co-culture with EL4 (Top) or E.G7-OVA (Bottom) cells. **f,** Cd8a surface expression of Cas9/OT-I cells transduced with Cd8a-targeting sgRNA (Red) compared to NC (Black) detected by flow cytometry. **g,** Normal distribution of sgRNAs in the sublibrary showing mean (Red) and median (Blue) values close to each other. **h,** Growth curve of tumors from E.G7-OVA cells inoculated s.c. into C57BL/6N mice (n=4). Data are shown as mean ± SEM. Black arrow indicates the time of sub-lethal irradiation followed by immediate adoptive transfer of Cas9/OT-I cells transduced with sgRNA sub-library.

**Extended Data Fig. 2.**
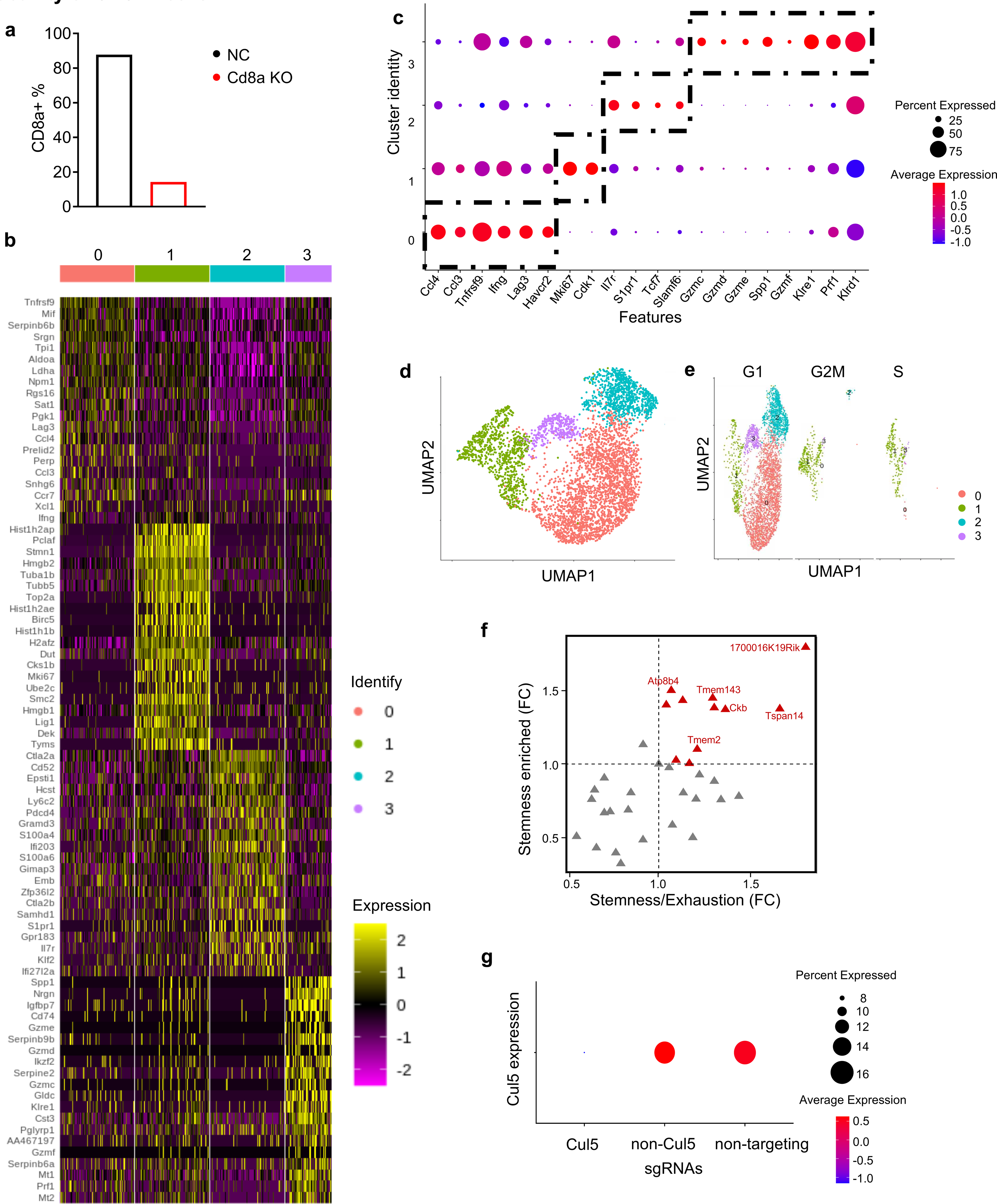
*In vivo* single-cell CRISPR KO screen identifies genes enhancing anti-tumor activity of CD8^+^ T cells. **a,** Flow cytometry analysis of CD8a expression on Cas9/OT-I T cells transduced with single-cell compatible non-targeting-(Black) or CD8a-sgRNA-containing (Red) retroviruses. **b,** Heatmap of differentially expressed genes in each cluster of tumor-infiltrating Cas9/OT-I T cells derived from single-cell RNA sequencing data. **c,** Dotplot of the expression of key DEGs in each cluster used to annotate T cell subtypes. **d,** UMAP projection of transferred tumor-infiltrating Cas9/OT-I cells transduced with the second sgRNA sub-library with 10x scaffold, colored by DEG clusters. **e,** UMAP visualization of cell clusters in cell cycle phases (G1, G2M and S). **f,** At gene level normalized to non-targeting control, sgRNA ratio of stemness cluster to the exhausted cluster as X axis and sgRNA enrichment in stemness cluster compared to input as Y axis. **g,** Dotplot of Cul5 gene expression in transferred tumor-infiltrating Cas9/OT-I cells expressing Cul5, non-targeting and the other sgRNAs.

**Extended Data Fig. 3.**
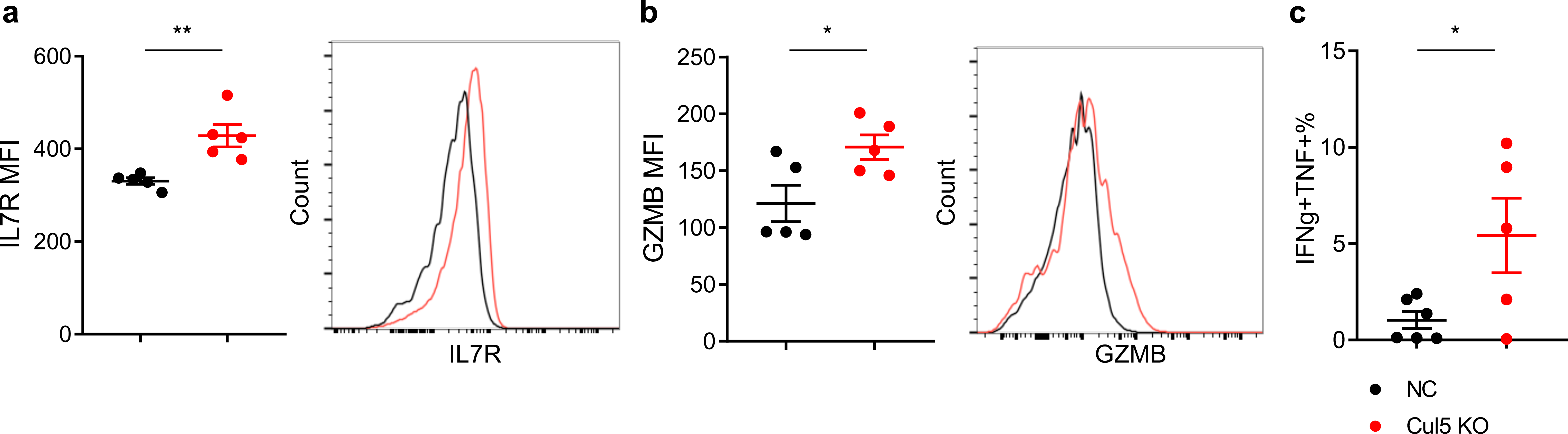
Cul5 KO enhances anti-tumor responses in primary CD8^+^ T cells *in vivo*. **a,b,** Flow cytometry analysis of **a,** CD127 and **b,** Granzyme B expression of transferred TDLN-infiltrating Cas9/OT-I cells with Cul5 or non-targeting KO. Left panel in each plot is a scatter plot of median fluorescent intensity (MFI) for each marker. Right panel is a representative histogram of each marker expression in Cul5 (Red) or NC (Black) KO T cells. Data are shown as mean ± SEM. *p < 0.05 and **p < 0.01, by unpaired t-test. **c,** Flow cytometry analysis of IFNg^+^TNF^+^ cell counts per unit tumor of transferred tumor-infiltrating Cas9/OT-I cells with Cul5 (Red) or NC (Black) KO post re-stimulation *in vitro*. Data are shown as mean ± SEM. *p < 0.05, by unpaired t-test. n=5-6

**Extended Data Fig. 4.**
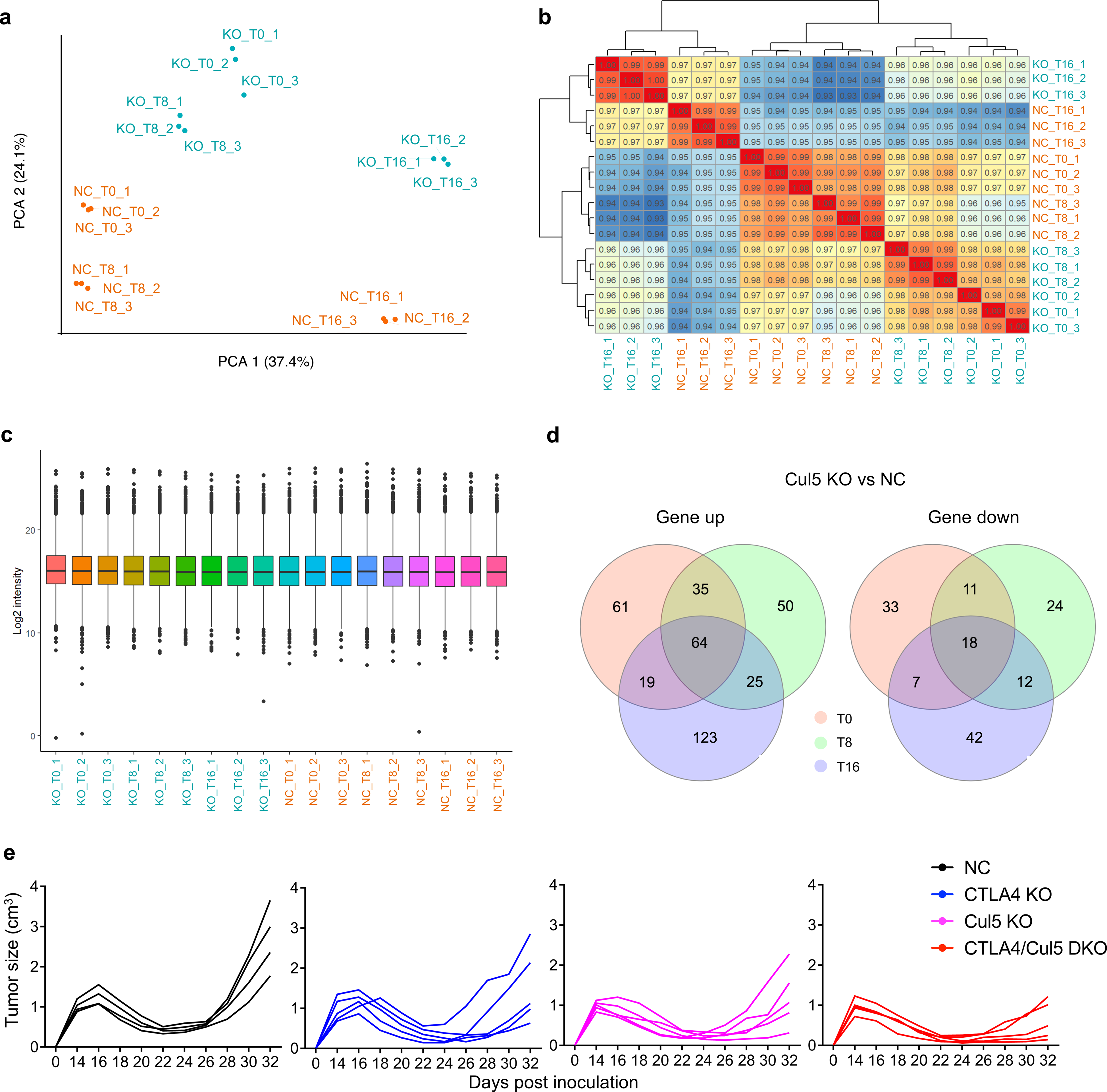
Cul5 KO causes proteomic alterations in primary CD8^+^ T cells. **a,** Principal component analysis of individual samples by the quantitative proteomics data using DIA-MS. **b,** Pearson correlation analysis between individual samples by proteomics data visualizing sample relationships. **c,** Normalized protein DIA-MS signals of individual samples by proteomics data. **d,** Venn diagram of proteins upregulated (Left) or downregulated (Right) in Cul5 KO primary CD8^+^ T cells compared to NC ones at T0, T8 and T16 conditions. **e,** Growth curve of tumors from E.G7-OVA cells inoculated s.c. into C57BL/6 mice. Data are shown as individual mouse. Black arrow indicates the time of sub-lethal irradiation followed by immediate adoptive transfer of Cas9/OT-I cells with NC (Black), Ctla4 (Blue), Cul5 (Pink) and Ctla4/Cul5 double (Red) KO. (n=4-5 mice per group).

**Extended Data Fig. 5.**
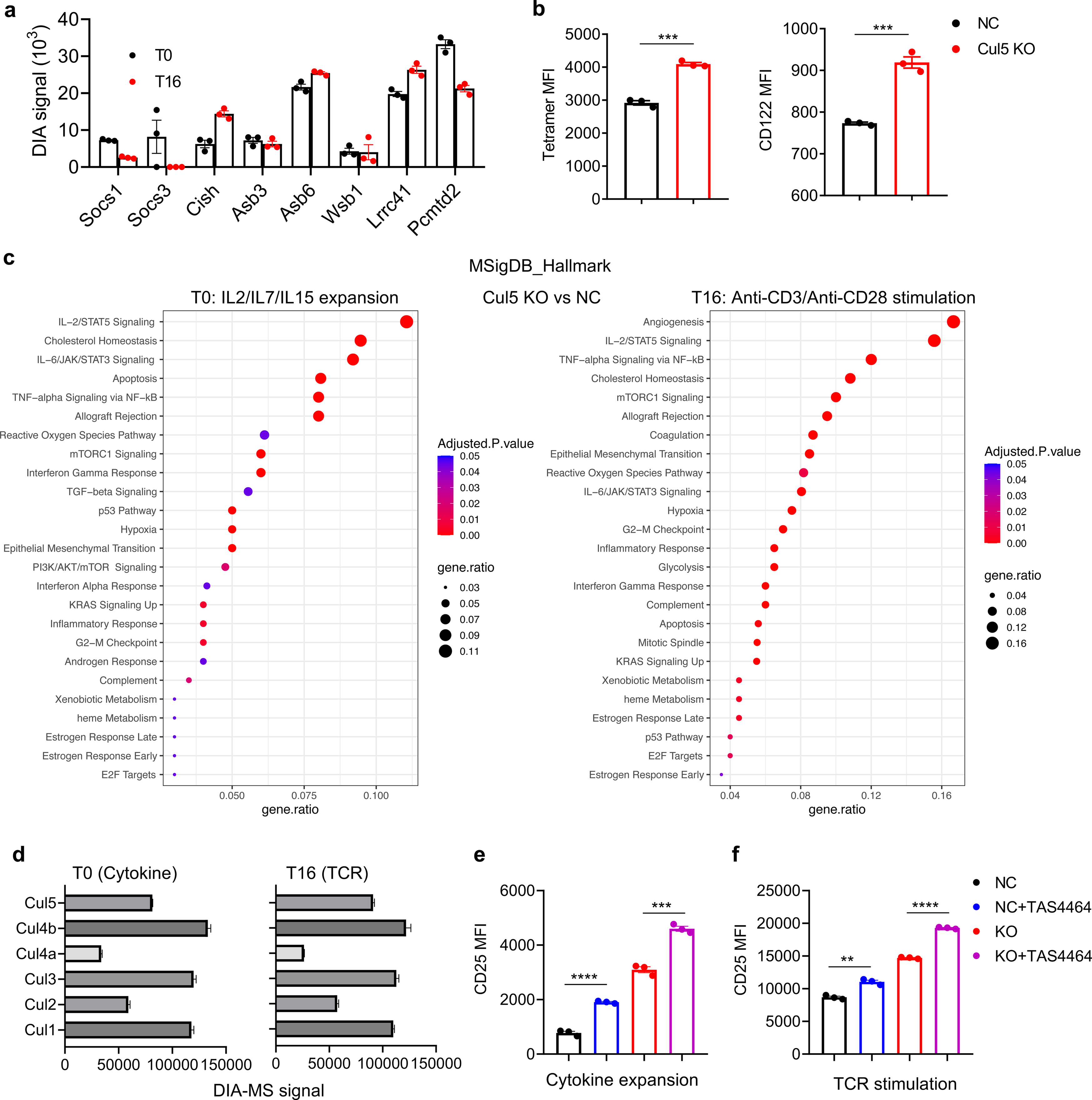
Signaling pathway enrichment and neddylation inhibition in CD8^+^ T cells with Cul5 KO. **a,** DIA-MS signals of different SOCS-box-containing proteins identified by the total protein MS analysis of NC mouse primary CD8^+^ T cells under T0 (cytokine expansion, Black) and T16 (TCR stimulation, Red) conditions. **b,** Flow cytometry analysis of TCR complex (SIINFEKL-H-2K(b) tetramer^+^, Left) and Il2rb (CD122, Right) expression of 16-hour anti-CD3/CD28-stimulated Cas9/OT-I cells with Cul5 (Red) or non-targeting (Black) KO. Data are shown as mean ± SEM. ***p < 0.001, by unpaired t-test, n=3. **c,** MSigDB Hallmark signaling pathway enrichment in Cul5 KO primary CD8^+^ T cells at T0 (Left) and T16 (Right) conditions. **d,** DIA-MS signals of different cullin proteins in NC mouse primary CD8^+^ T cells under T0 (cytokine expansion) and T16 (TCR stimulation) conditions. **e,f,** Flow cytometry analysis of CD25 in NC (Black and Blue) or Cul5 KO (Red and Purple) primary CD8^+^ T cells in 16-hour **e,** cytokine culture (IL2/IL7/IL15) or **f,** anti-CD3 plus anti-CD28 stimulation with (Blue and Purple) or without (Black and Red) 250nM TAS4464 addition. Data are shown as mean ± SEM. **p < 0.01, ***p < 0.001, and ****p < 0.0001 by unpaired t-test, n=3.

**Extended Data Fig. 6.**
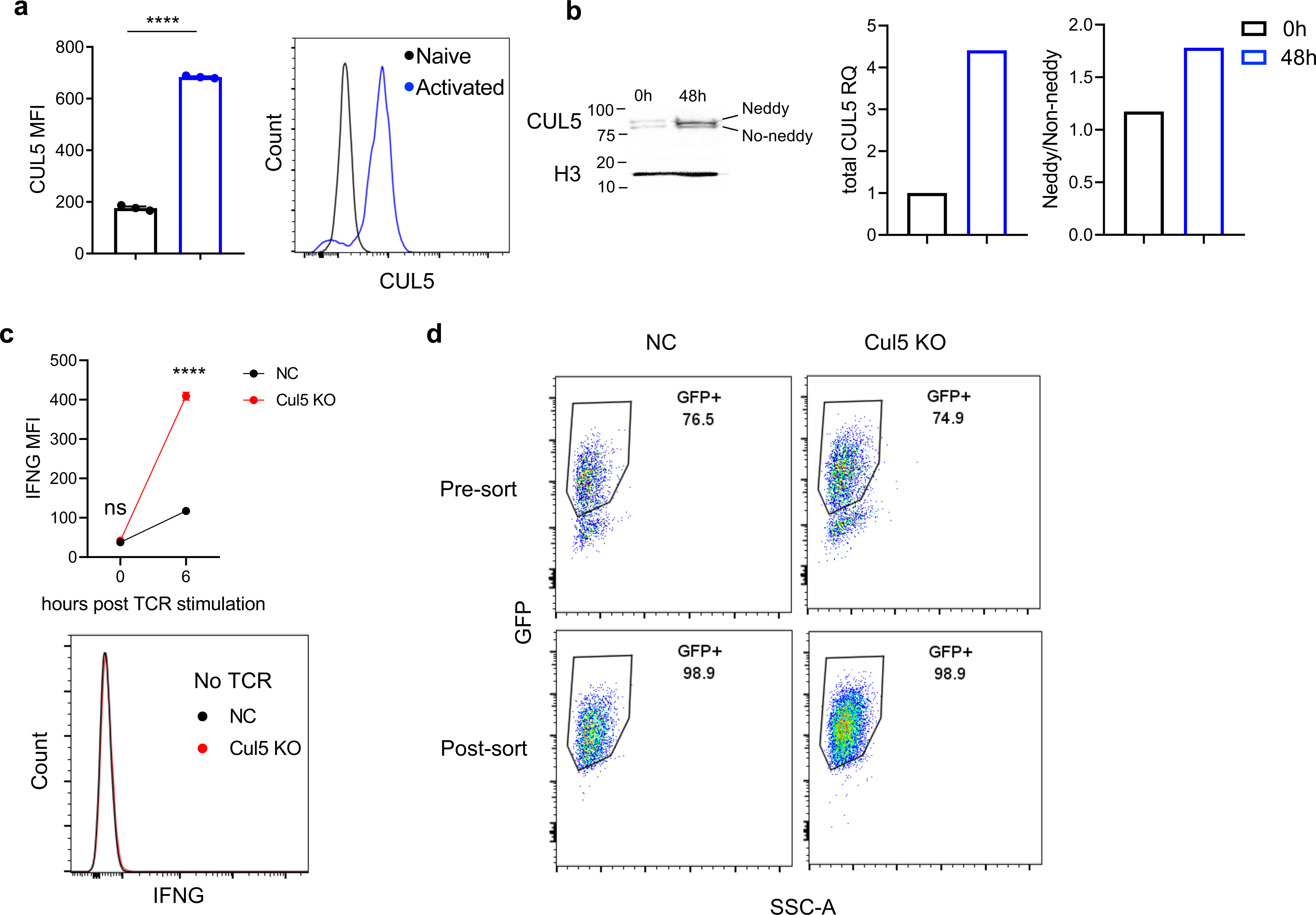
CUL5 KO in primary human CD8^+^ T cells. **a,** Flow cytometry analysis of CUL5 expression in primary human CD8^+^ T cells before (Black) or after (Blue) 2-day activation by anti-CD3/CD28 beads. Data are shown as mean ± SEM (****p < 0.0001, by unpaired t-test, n=3). **b,** Western blot analysis of CUL5 expression in primary human CD8^+^ T cells treated as in **a**. Histone H3 as an internal control. Bands were shown in the left panel. Relative quantity (RQ) of total CUL5 protein was normalized to Histone H3 in the middle panel. The ratio of neddylated to non-neddylated CUL5 was shown in the right panel. **c,** Flow cytometry analysis of IFNG MFI in NC (Black) or CUL5 KO (Red) primary human CD8^+^ T cells before and after 6-hour anti-CD3 plus anti-CD28 stimulation. Data are shown as mean ± SEM (ns as not significant and ****p<0.0001 by unpaired t-test, n=3). **d,** Flow cytometry analysis of human CD8^+^ CAR-CD19-T cells with NC or CUL5 KO pre- or post-sorting of GFP^+^ population.

**Extended Data Fig. 7.**
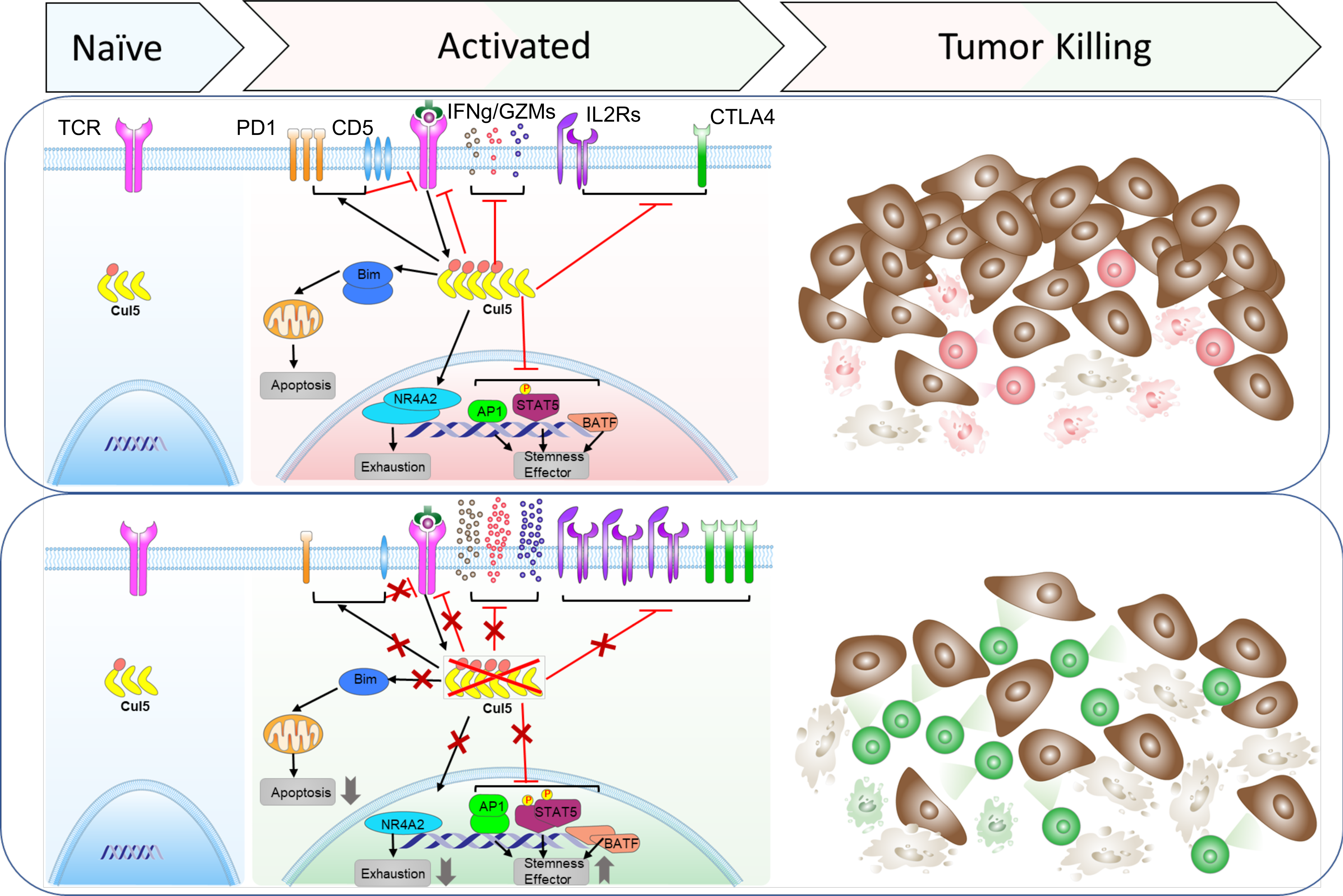
Illustration summary of Cul5 regulation of CD8^+^ T cell anti-tumor responses.

